# Alveolar macrophages strictly rely on GM-CSF from alveolar epithelial type 2 cells before and after birth

**DOI:** 10.1101/2021.04.01.438051

**Authors:** Julia Gschwend, Samantha Sherman, Frederike Ridder, Xiaogang Feng, Hong-Erh Liang, Richard M. Locksley, Burkhard Becher, Christoph Schneider

## Abstract

Programs defining tissue-resident macrophage identity depend on local environmental cues. For alveolar macrophages (AMs), these signals are provided by immune and non-immune cells, and include GM-CSF (CSF2). However, evidence to functionally link components of this intercellular crosstalk remains scarce. We thus developed new transgenic mice to profile pulmonary GM-CSF expression, which we detected in both immune cells, including group 2 innate lymphoid cells and γδ T cells, as well as AT2s. AMs were unaffected by constitutive deletion of hematopoietic *Csf2* and basophil depletion. Instead, AT2 lineage-specific constitutive and inducible *Csf2* deletion revealed the non-redundant function of AT2-derived GM-CSF in instructing AM fate, establishing the postnatal AM compartment, and maintaining AMs in adult lungs. This AT2-AM relationship begins during embryogenesis, where nascent AT2s timely induce GM-CSF expression to support the proliferation and differentiation of fetal monocytes contemporaneously seeding the tissue, and persists into adulthood, when epithelial GM-CSF remains restricted to AT2s.

## INTRODUCTION

Proper organ development and homeostasis is reliant not only on the appropriate function of specialized tissue-specific stromal cells, but also on the crosstalk that occurs between these cells and the stably or transiently integrated tissue-resident immune cell populations. Tissue-resident macrophages (TRMs) are often integral members of these immune cross-talk compartments, and are best known for their phagocytic activity (Davies et al., 2013). Work over the past years has identified distinct transcriptional programs that are associated with particular tissue-specific functions in TRM subsets (Blériot et al., 2020). Instigation of these programs is particularly critical during the embryonic development of an organ, when TRMs first differentiate from different myeloid progenitors and readily acquire tissue-specific transcriptional signatures (Mass et al., 2016). Although the processes of TRM induction are not well defined, it is presumed that these programs are initiated and maintained by signals derived from the local tissue microenvironment (Guilliams et al., 2020).

Alveolar macrophages (AMs) are a specific type of TRM in the lung. They play a central role in maintenance of alveolar homeostasis by removing cellular debris, excess surfactant and inhaled bacterial, but are also important in preserving lung function during pulmonary viral infections (Hussell and Bell, 2014; Kopf et al., 2015). Found within the lumen of the alveoli, AMs are surrounded by two alveolar epithelial cell types. Alveolar type 1 cells (AT1s) facilitate gas exchange with underlying endothelial cells, while alveolar type 2 cells (AT2s) produce surfactant, act as facultative progenitors in case of alveolar injury, and activate the immune system in response to pathogen-related stimuli (Basil et al., 2020; Whitsett et al., 2019). Notably, this epithelial niche is formed towards the end of embryonic lung development, and its establishment is concomitant with the differentiation of AMs (Guilliams et al., 2013; Schneider et al., 2014b). Work generated over the past several years suggests that differentiating AMs pass through a transitional monocytic stage during their development (Evren et al., 2021; Guilliams et al., 2013; Hoeffel et al., 2015; Liu et al., 2019; Schneider et al., 2014b; Yona et al., 2013), while parabiosis and fate-mapping studies revealed that, similar to other TRM compartments, AMs comprise cells of both fetal and adult monocytic origin (Hashimoto et al., 2013; Schulz et al., 2012; Yona et al., 2013). Novel refined tools enabled the identification of multiple myeloid progenitors which contribute to the pool of AMs and which are produced during partially overlapping successive waves of hematopoiesis (Gomez Perdiguero et al., 2015; Hoeffel et al., 2015; Liu et al., 2019). Notably, experiments using mixed bone marrow chimeras and adoptive transfers of fetal and adult precursor populations demonstrated that regardless of origin, a major determinant for the generation of a functional AM compartment in mice and humans is signaling via the cytokine granulocyte-macrophage colony-stimulating factor (GM-CSF, encoded by the gene *Csf2*) (Guilliams et al., 2013; Li et al., 2020; Schneider et al., 2017; Schneider et al., 2014a; Schneider et al., 2014b; Suzuki et al., 2014; van de Laar et al., 2016). Indeed, more than 30 years ago, Chen et al. speculated about GM-CSF-mediated AM self-renewal, which is decoupled from bone marrow hematopoiesis (Chen et al., 1988). The critical function of GM-CSF for AMs and surfactant homeostasis was later confirmed in knockout mice with deficiencies in the GM-CSF signaling pathway (Dranoff et al., 1994; Nishinakamura et al., 1995; Robb et al., 1995; Stanley et al., 1994). Subsequent findings in patients with rare *Csf2* mutations or anti-GM-CSF autoantibodies and in studies with humanized mice demonstrated that this distinctive lung GM-CSF function is fundamentally conserved in humans (Kitamura et al., 1999; Martinez-Moczygemba et al., 2008; Suzuki et al., 2008; Willinger et al., 2011). Mouse studies have demonstrated that the binding of GM-CSF to its receptor (encoded by *Csf2ra* and *Csf2rb*) is associated with induction of the transcription factor peroxisome proliferator-activated receptor gamma (PPARγ), and the corresponding initiation of AM differentiation in lung fetal monocytes during prenatal development (Schneider et al., 2014b). PPARγ is thought to critically regulate an AM-specific transcriptional program, including upregulation of signature markers CD11c and Siglec-F, and to drive terminal AM differentiation in the first 1-2 weeks after birth.

Disruption of the mature functional AM population compromises lung homeostasis and in both humans and mice. Loss of GM-CSF signaling and the subsequent deficit of AMs leads to the accumulation of lipoproteinaceous material and cellular debris in the alveolar space, a condition known as pulmonary alveolar proteinosis (PAP) (Trapnell et al., 2019). These detrimental phenotypes can be rescued in mice by re-establishing a functional AM compartment via pulmonary overexpression of GM-CSF in *Csf2*^−/−^ mice (Huffman et al., 1996), or by supplying *Csf2r*-deficient mice with cells that have both intact *Csf2r* and the potential to differentiate into mature AMs (Schneider et al., 2014a; Suzuki et al., 2014). Thus, in addition to its known function in conferring an inflammatory state in myeloid cells under inflammatory and autoimmune conditions (Becher et al., 2016; Hamilton, 2020), GM-CSF also has a dedicated role in shaping lung AM fate (Evren et al., 2020). Indeed, results from early studies published almost 50 years ago using mouse lung conditioned medium indicated high GM-CSF expression in the lung (Sheridan and Metcalf, 1973); the responsible cells, however, remained elusive. Production of GM-CSF in the lung has since been attributed to multiple hematopoietic and non-hematopoietic lineages, including type 2 innate lymphoid cells (ILC2s), basophils, and epithelial cells, specifically AT2s (Cohen et al., 2018; Guilliams et al., 2013; Schneider et al., 2014b). Basophils and ILC2s were suggested to constitute an important part of this AM niche and to be critical regulators of AM differentiation and function (Cohen et al., 2018). Other sources of GM-CSF may include T cells, B cells, natural killer (NK) cells, ILCs, fibroblasts, and endothelial cells, as suggested by data obtained from different tissues under inflammatory conditions (Hamilton, 2020). However, it is unclear which cellular sources of GM-CSF are most critical for promoting AM differentiation. It is also not known whether the critical GM-CSF producers differ between embryonic, neonatal and adult lungs, nor if this axis regulates AM maintenance following perinatal AM establishment.

To unambiguously identify the relevant sources of GM-CSF required the formation of AMs *in vivo*, we thus generated novel GM-CSF reporter and conditional knockout mice, and proceeded to systematically delete GM-CSF from different hematopoietic and non-hematopoietic compartments. Here, we identify AT2s as a constitutive and dominant non-hematopoietic source of GM-CSF in the lungs of young and adult mice. We further demonstrate that AT2-derived GM-CSF is the necessary signal to initiate and promote pre- and post-natal AM differentiation and subsequent AM maintenance throughout adulthood. GM-CSF is a lineage-defining cytokine in lung epithelial cells, which becomes expressed following AT2 fate specification during late gestation contemporaneously with AM fate initiation in lung fetal monocytes. Our results unequivocally demonstrate that AMs are critically reliant not on hematopoietic GM-CSF but rather on epithelial GM-CSF derived from AT2s, both for development and for maintenance of the mature AM population.

## RESULTS

### ILC2s are the major hematopoietic source of *Csf2* expression in the lungs

To disentangle the individual contributions of different pulmonary GM-CSF sources, we generated a knock-in mouse model that was designed to report *Csf2* expression and to allow for *Csf2* deletion (**Fig. 1A**). In the absence of Cre, transcription of the *Csf2*^flox-tdTomato^ locus (*Csf2*^fl^) results in the production of a bicistronic mRNA encoding GM-CSF and tdTomato, marking GM-CSF producing cells with fluorescent tdTomato signal while maintaining normal GM-CSF expression. When recombined in the germline to create mice with a global deletion of *Csf2* (*Csf2*^Δ^), we found that AMs were completely absent from the adult lung, as per other *Csf2*^−/−^ models; these results validated the loss-of-function property of this novel mouse tool (**Fig. 1B**). Evaluation of the reporter function of this tool revealed that tdTomato^+^ cells were readily detectable in the lungs of *Csf2*^+/fl^ mice compared to *Csf2*^+/+^ littermate controls (**Fig. 1C**). Notably, the results suggested the presence of multiple GM-CSF sources, including cells of hematopoietic (CD45^+^) and non-hematopoietic (CD45^−^) origin (**Fig. 1C**). Furthermore, when the floxed allele was deleted (*Csf2^+/^*^Δ^), this resulted in strongly reduced tdTomato expression in both compartments (**Fig. 1C**). This unexpected loss of tdTomato signal in the case of *Csf2*^Δ^ alleles, however, provided us with an endogenous readout for assessing Cre-mediated *Csf2* deletion.

**Figure 1.**
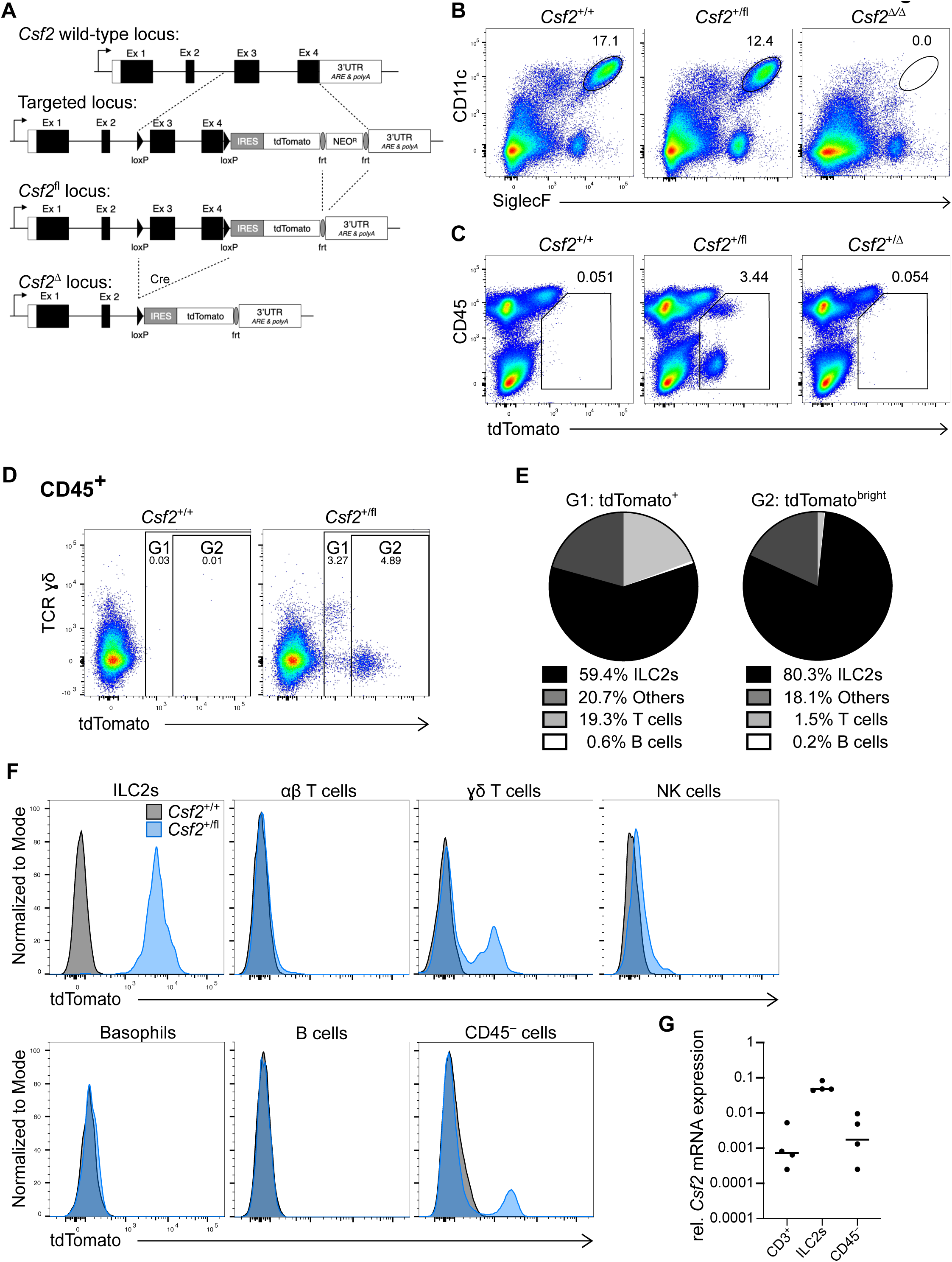
Relative GM-CSF contribution of different hematopoietic cells in the neonatal lung. (**A**) Gene-targeting strategy used to engineer the *Csf2*^flox-tdTomato^ (*Csf2*^fl^) mice. (**B**) Flow cytometry analysis of CD11c^+^SiglecF^+^ AMs in the lungs of adult *Csf2*^+/+^, *Csf2*^+/fl^ and *Csf2*^Δ/Δ^ mice, gated on live CD45^+^ cells. (**C**) Flow cytometry analysis of tdTomato^+^ populations in the adult lungs of *Csf2*^+/+^, *Csf2*^+/fl^ and *Csf2*^Δ/fl^ mice, gated on live cells. (**D**) Flow cytometry of tdTomato expression by CD45^+^ cells in *Csf2*^+/+^ and *Csf2*^+/fl^ P10 lungs, gated on live cells. (**E**) Percentage contribution from different hematopoietic cells types to the CD45^+^tdTomato^+^ (left) or CD45^+^tdTomato^high^ compartment, as gated in E. (**F**) tdTomato signal from the indicated cell populations in the lungs of P10 *Csf2*^+/+^ (grey) and *Csf2*^+/fl^ (blue) mice. Expression of tdTomato was determined by flow cytometry analysis. (**G**) *Csf2* mRNA expression relative to *Rps17* in the three major tdTomato^+^ cell populations (ILC2s, T cells and CD45^−^ cells). Cells were sorted from P10 *Csf2*^+/fl^ lungs. (**B+C**) Data are from one experiment representative of two independent experiments. (**D-F**) Data are from one experiment representative of five independent experiments. (**G**) Pooled data from three independent experiments.

Because AMs complete their development shortly after birth (Guilliams et al., 2013; Schneider et al., 2014b), we first characterized the major sources of GM-CSF in the neonatal lung. For this, postnatal day 10 (P10) lungs from *Csf2*^+/+^ and *Csf2*^+/fl^ mice were analyzed by flow cytometry. As observed in the adult lungs (**Fig. 1C**), the tdTomato^+^ cells from neonatal *Csf2*^+/fl^ mice were comprised of CD45^−^ and CD45^+^ populations, of which the latter showed slightly higher tdTomato expression (**Supp. Fig. 1A**). Further analysis of the CD45^+^tdTomato^+^ cells identified several different cell types, including Lin^−^Thy1^+^ST2^+^ ILC2s, CD3^+^TCRβ^+^ T cells, and CD3^+^TCRγδ^+^ T cells (**Supp. Fig. 1B**). ILC2s were found to be the major hematopoietic GM-CSF producers by frequency (> 80%) and by relative reporter expression, followed by CD3^+^ T cells; other minor contributions to the tdTomato signal came from NK cells and B cells (**Fig. 1D-F, Supp. Fig. 1B**). ILC2s produced a strong reporter signal, with nearly all of the *Csf2*^+/fl^ ILC2s expressing tdTomato (**Fig. 1F**). In contrast, very few of the T cells expressed tdTomato, and the overall tdTomato fluorescent signal was not nearly as strong as was observed with the ILC2s (**Fig. 1F**). To further confirm that the tdTomato signal was reporting *Csf2* expression, qRT-PCR was used to compare the relative *Csf2* mRNA in sorted tdTomato^+^ ILC2s, tdTomato^+^CD3^+^ cells, and tdTomato^+^CD45^−^ cells. Overall, higher levels of *Csf2* mRNA were detected in the ILC2s relative to T cells and CD45^−^ cells (**Fig. 1G**). Together these results suggest that in the neonatal lung, ILC2s express the overall highest amount of GM-CSF on a per-cell basis and also represent the dominant GM-CSF source within the hematopoietic compartment, which is in agreement with a prior report (Cohen et al., 2018).

### Hematopoietic-derived GM-CSF is dispensable for the development of AMs in the neonatal lung

Multiple studies suggest that ILC2s shape the neonatal lung environment via their canonical type 2 effector cytokines, IL-5 and IL-13 (Cohen et al., 2018; de Kleer et al., 2016; Nussbaum et al., 2013; Saluzzo et al., 2017; Schneider et al., 2019; Steer et al., 2017). Indeed, the constitutive *Csf2* expression by postnatal lung ILC2s is reminiscent of their production of IL-5, by which they regulate eosinophil homeostasis (Nussbaum et al., 2013). To determine the relative importance of hematopoietic GM-CSF for AM homeostasis, including GM-CSF derived from ILC2s, we deleted *Csf2* expression from the hematopoietic compartment by crossing *Csf2*^fl^ mice with *Vav1*^iCre^ mice and analyzed P10 neonatal lungs by flow cytometry. For technical reasons and due to occasional germline Cre activity in *Vav1*^iCre^ mice (see methods section), experimental cohorts included mice with *Csf2*^Δ/fl^ and *Csf2*^fl/fl^ alleles (hereafter termed ‘*Csf2*^fl^’). All conditional KO groups were compared to their littermate Cre^−^ controls. In comparison to Cre^−^ controls, analysis of P10 lungs from *Vav1*^iCre^;*Csf2*^fl^ mice revealed a reduction in tdTomato signal in CD45^+^ cells, while the tdTomato signal from the CD45^−^ compartment was unchanged (**Fig. 2A, B**). Absence of GM-CSF protein expression in ILC2s from *Vav1*^iCre^;*Csf2*^fl^ mice was further validated by *in vitro* restimulation of lung cells (**Fig. 2C, D**). Following confirmation of specific and efficient depletion of hematopoietic-derived GM-CSF in *Vav1*^iCre^;*Csf2*^fl^ mice, their myeloid compartment was analyzed for AMs. Assessment of the AM population indicated that there was no change in the number of AMs in the lung in the absence of hematopoietic *Csf2* expression (**Fig. 2E, F**).

**Figure 2.**
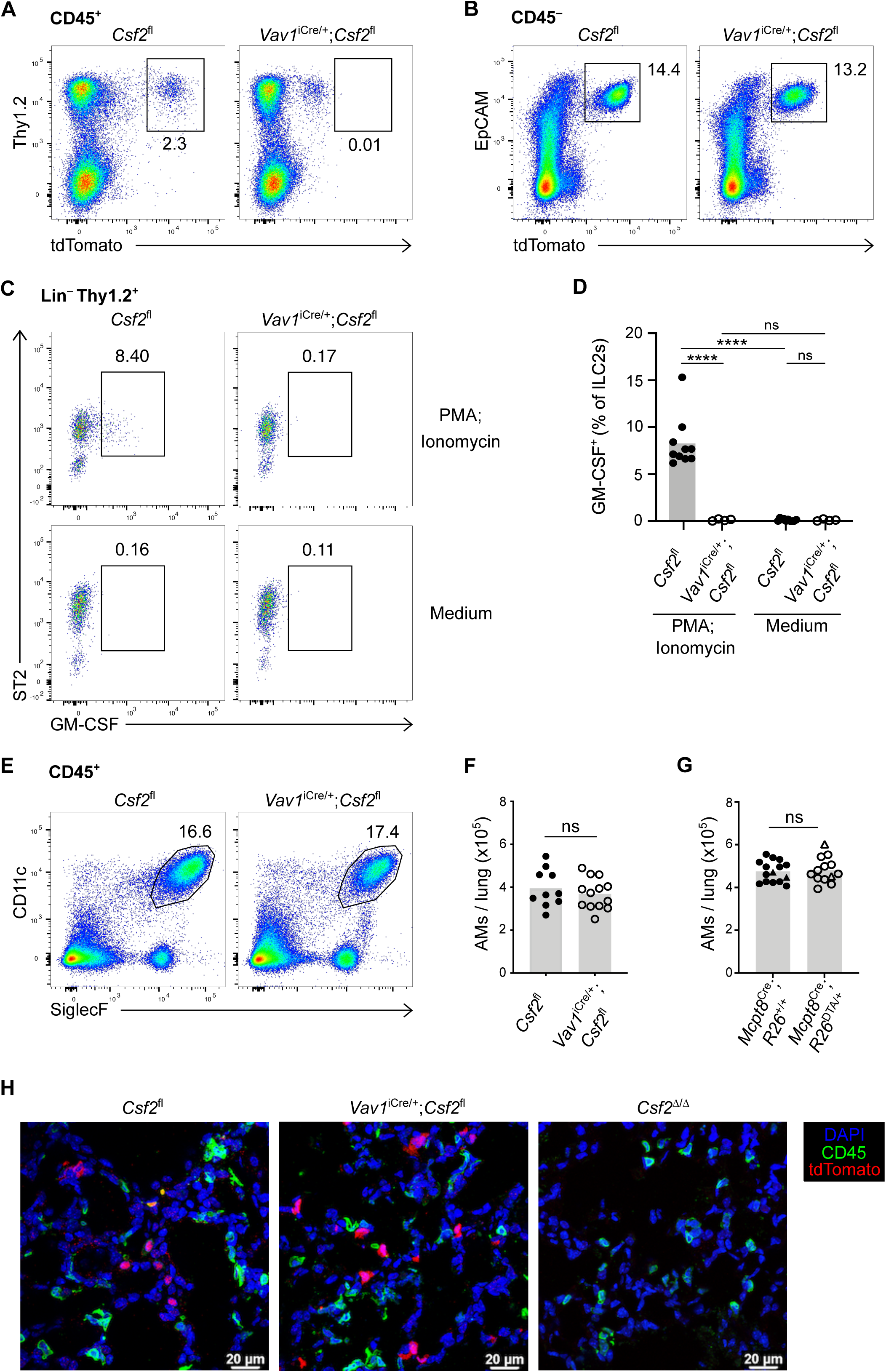
Hematopoietic-derived GM-CSF is dispensable for the development of AMs in the neonatal lung. (**A**) Flow cytometry analysis of CD45^+^tdTomato^+^ populations in P10 lungs of *Csf2*^fl^ and *Vav1*^iCre/+^;*Csf2*^fl^ mice, gated on live cells. (**B**) Flow cytometry analysis of CD45^−^tdTomato^+^ populations in P10 lungs of *Csf2*^Fl^ and *Vav1*^iCre/+^;*Csf2*^fl^ mice, gated on live cells. (**C**) Flow cytometry analysis of GM-CSF production in ILC2s (CD45^+^Lin^−^Thy1.2^+^ST2^+^) from P10 lungs of *Csf2*^fl^ and *Vav1*^iCre/+^;*Csf2*^fl^ mice after restimulation with PMA/ionomycin (top row) or incubation with medium only (bottom row). (**D**) Percentage of GM-CSF^+^ ILC2s in P10 lungs of *Csf2*^fl^ and *Vav1*^iCre/+^;*Csf2*^fl^ mice after restimulation or in the presence of medium only. (**E**) Flow cytometry analysis of CD11c^+^SiglecF^+^ AMs in P10 lungs of *Csf2*^fl^ and *Vav1*^iCre/+^;*Csf2*^fl^ mice, gated on live CD45^+^ cells. (**F**) AM (CD45^+^Ly-6G^−^SiglecF^+^CD11c^+^CD64^+^) quantification in P10 lungs of *Csf2*^fl^ and *Vav1*^iCre/+^;*Csf2*^fl^ mice. (**G**) AM (CD45^+^SiglecF^+^CD11c^+^) quantification in P10 lungs of *Mcpt8*^YFP-Cre^;*R26*^+/+^ and *Mcpt8*^YFP-Cre^;*R26*^DTA/+^ mice. *Mcpt8*^YFP-Cre/+^ mice are indicated by circles, while *Mcpt8*^YFP-Cre/Cre^ mice are indicated by triangles. (**H**) Representative IF pictures of tdTomato^+^ cells (red) and CD45^+^ cells (green) in P10 lungs of *Csf2*^fl^, *Vav1*^iCre/+^;*Csf2*^fl^ and *Csf2*^Δ/Δ^ mice. (**A, B, E, H**) Data are from one experiment representative of at least two independent experiments. (**C**) Data are from one experiment, representative of two independent experiments. (**D**) Data are pooled from two independent experiments. (**F**) Data pooled from four independent experiments. **(G)** Data are pooled from three independent experiments.

A previous report, however, suggests that basophils are important regulators of AM development, a function that was in part attributed to the GM-CSF produced by ILC2s and/or basophils themselves (Cohen et al., 2018). Since our results did not support a critical contribution of hematopoietic-derived GM-CSF to AM survival, we further explored the role of basophils in this process by using a validated genetic basophil depletion model (Sullivan et al., 2011), in which basophil-specific Cre expression (*Mcpt8*^YFP-Cre^) ablates YFP-tagged basophils through *Rosa26*-driven expression of the cell-lethal diphtheria toxin A (DTA). The constitutive depletion of basophils in *Mcpt8*^YFP-Cre^;*R26*^DTA^ mice (**Supp. Fig. 2A, B**), however, did not significantly alter AM numbers in the P10 lung even when using two different gating strategies (**Fig. 2G, Supp. Fig. 2C-E**). Furthermore, we did not detect changes in their canonical surface marker expression when compared to AMs from littermate control mice (**Supp. Fig. 2F, G**).

Overall, these results demonstrate that neither the constitutive loss of hematopoietic *Csf2* expression nor the depletion of basophils *per se* affects the number of AMs in the developing neonatal lung. Indeed, despite a substantial reduction in the *Csf2*-reporter signal in dissociated lungs from *Vav1*^iCre^;*Csf2*^fl^ mice, *in situ* analysis by immunofluorescence revealed that the overall tdTomato signal was still present in these lungs (**Fig. 2H**). Furthermore, the lack of CD45 co-expression in the remaining tdTomato^+^ cells suggests that these cells exclusively belong to the CD45^−^ compartment, and are more numerous than initially anticipated from flow cytometry analysis of dissociated tissue. These studies thus point towards the CD45^−^ compartment as the major orchestrator of GM-CSF-mediated AM differentiation and survival.

### Epithelial AT2s are the main source of *Csf2* expression in the non-hematopoietic compartment of the neonatal lung

To identify the CD45^−^ cell types that express the *Csf2*-reporter, neonatal P10 lungs from *Csf2*^+/+^ and *Csf2*^+/fl^ mice were profiled by flow cytometry using a comprehensive panel of markers for epithelial, mesenchymal, and endothelial lineages. tdTomato expression was only detected in EpCAM^+^ epithelial cells that co-expressed MHCII; no *Csf2*-reporter signal was detected in CD45^−^EpCAM^−^ cells, including CD31^+^ endothelial and PDGFRɑ^+^ mesenchymal populations (**Fig. 3A**). Airway epithelial cells (EpCAM^+^CD104^+^) were also found to be tdTomato^−^. Further characterization of the tdTomato^+^ epithelial cells revealed that these cells also expressed pro-surfactant protein C (SP-C), and the sodium-dependent phosphate transporter NaPi-IIb (Slc34a2) (**Fig. 3B**). In the neonatal and adult lung, all three of these markers – MHCII, SPC, and NaPi-IIb – are very strongly associated with AT2s (Donati et al., 2020; Tighe et al., 2019; Traebert et al., 1999). Taken together, we conclude that AT2s constitutively express *Csf2* and that the vast majority of *Csf2*-expressing epithelial cells are AT2s, at least as assessed using this reporter in dissociated tissue of the neonatal lung.

**Figure 3.**
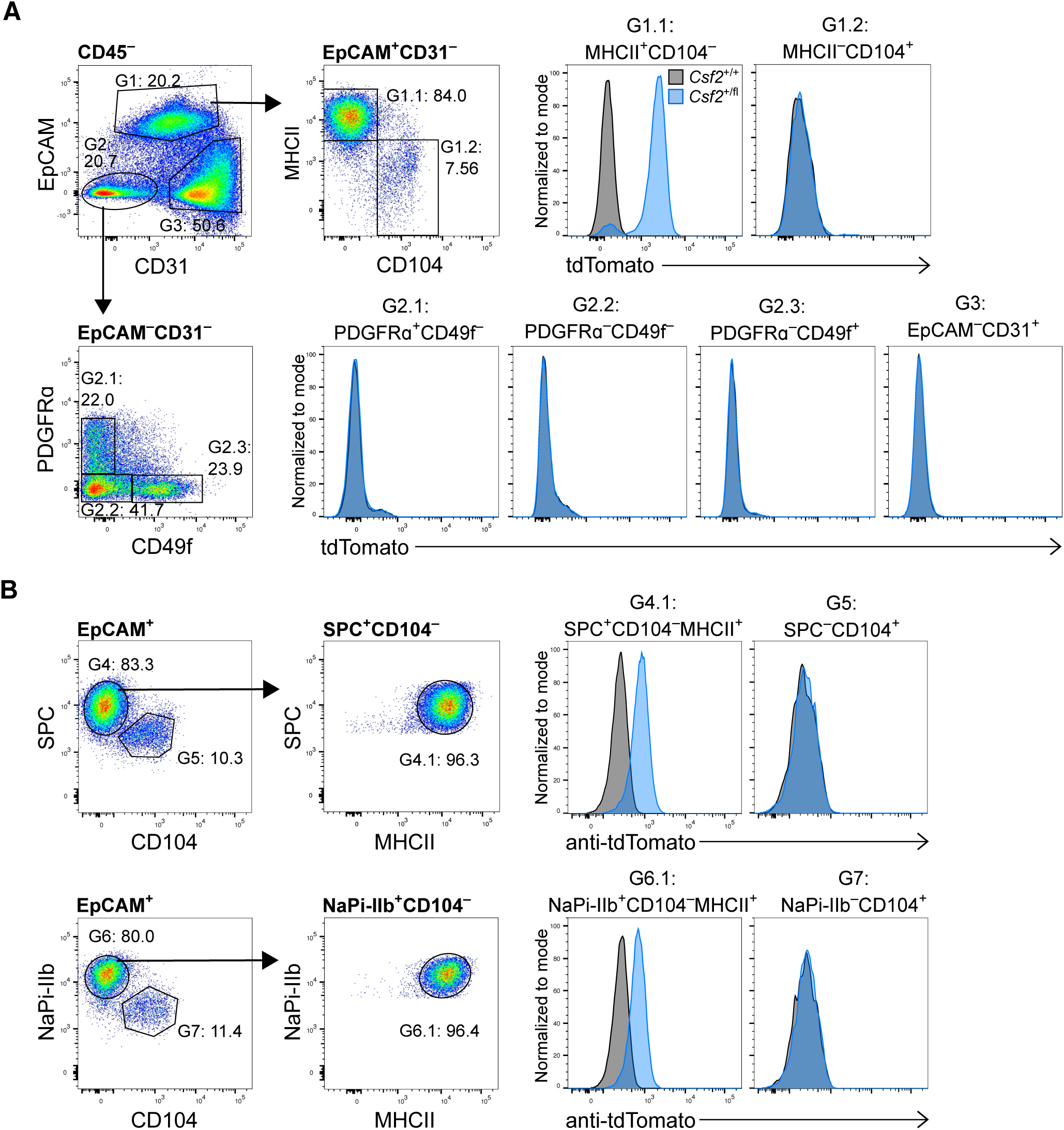
AT2s as are the main non-hematopoietic source of GM-CSF in the neonatal lung. (**A**) Flow cytometry analysis of tdTomato expression in different CD45^−^ cell populations in P10 lungs of *Csf2*^+/+^ (grey) and *Csf2*^+/fl^ (blue) mice. Isolated populations include AT2s (G1.1: CD45^−^CD31^−^EpCAM^+^MHCII^+^CD104^−^); airway epithelial cells (G1.2: CD45^−^CD31^−^EpCAM^+^MHCII^−^CD104^+^); fibroblasts (G2.1: CD45^−^CD31^−^ EpCAM^−^PDGFRɑ^+^CD49f^−^); G2.2: CD45^−^CD31^−^EpCAM^−^PDGFRɑ^−^CD49f^−^ cells; G2.3: CD45^−^CD31^−^ EpCAM^−^PDGFRɑ^−^CD49f^+^ cells; and endothelial cells (G3: CD45^−^CD31^+^EpCAM^−^). (**B**) Flow cytometry analysis of tdTomato expression in the CD45^−^CD31^−^EpCAM^+^ compartment in fixed P10 lungs of *Csf2*^+/+^ (grey) and *Csf2*^+/fl^ mice (blue) to further verify AT2 identity. Isolated populations include AT2s (G4.1: SPC^+^CD104^−^MHCII^+^; G6.1: NaPi-IIb^+^CD104^−^MHCII^+^) and airway epithelial cells (G5: SPC^−^CD104^+^; G7: NaPi-IIb^−^CD104^+^). (**A+B**) Data are from one experiment representative of two independent experiments.

### Depletion of AT2-derived *Csf2* expression abrogates AM development

Overall, our data suggest that the major production of GM-CSF in the neonatal lung can be specifically attributed to the AT2s. To test the functional relevance of AT2-derived GM-CSF for perinatal AM development, we generated *SPC*^Cre^;*Csf2*^fl^ mice, in which Cre expression is under the control of the human surfactant protein C gene promoter (Okubo and Hogan, 2004). Compared to the abundant tdTomato^+^ cells that were found in lung sections of *Csf2*^fl^ mice on P10, few tdTomato^+^ cells were detected in lungs of *SPC*^Cre^;*Csf2*^fl^ mice (**Fig. 4A**). Notably, these remaining tdTomato^+^ cells also expressed CD45, identifying them as hematopoietic cells. As expected, tdTomato signal was completely absent in *Csf2*^Δ/Δ^ lung sections (**Fig. 4A**). Flow cytometry analysis of dissociated lungs revealed that *Csf2*-reporter expression by CD45^−^ EpCAM^+^MHCII^+^ AT2s was uniformly reduced, whereas no change was detected in tdTomato expression by the CD45^+^ hematopoietic compartment nor, more specifically, the ILC2s (**Fig. 4B, C, Supp. Fig. 3A-C**). This suggests that the *SPC*^Cre^ was effective in specifically depleting *Csf2* expression in AT2s while leaving hematopoietic *Csf2* expression intact. These findings are supported by measurements of total GM-CSF in media conditioned with P10 lungs, in which GM-CSF levels in conditioned media from *SPC*^Cre^;*Csf2*^fl^ and *Csf2*^Δ/Δ^ lungs were significantly reduced compared to control *Csf2*^fl^ and *Vav1*^iCre^;*Csf2*^fl^ media samples (**Fig. 4D**). The functional consequences of deleting GM-CSF specifically from AT2s were then evaluated by examining the AM compartment in *SPC*^Cre^;*Csf2*^fl^ mice and their littermate controls. Remarkably, CD11c^+^SiglecF^+^ AMs were completely absent in *SPC*^Cre^;*Csf2*^fl^ mice (**Fig. 4E, F**). Overall, these results unambiguously demonstrate that *Csf2* expression from the epithelial compartment of the neonatal lung, specifically from AT2s, is crucial for AM differentiation and survival.

**Figure 4.**
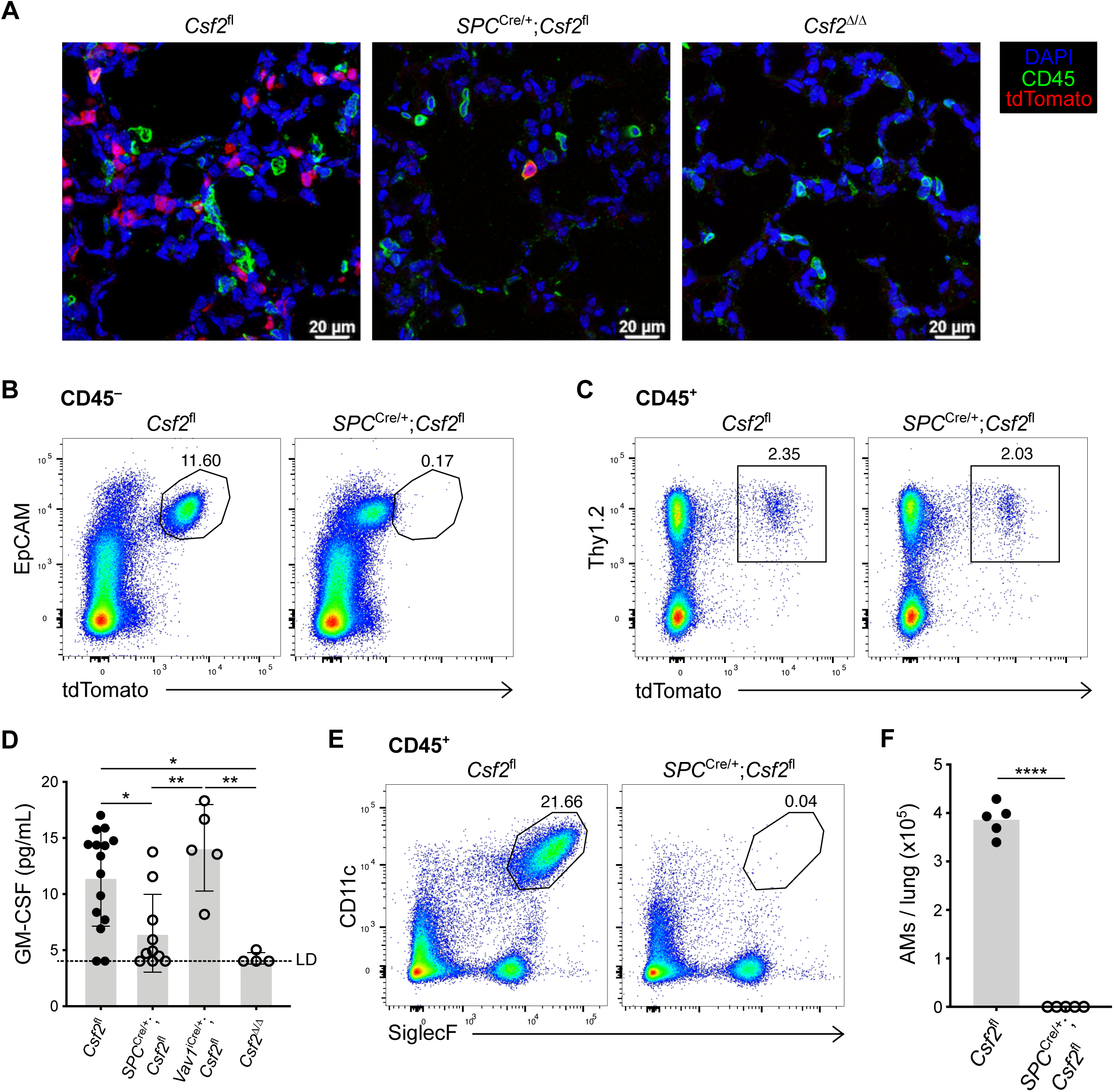
AT2-derived GM-CSF is necessary for the development of AMs in the neonatal lung. (**A**) Representative IF pictures of tdTomato^+^ cells (red) and CD45^+^ cells (green) in P10 lungs of *Csf2*^fl^, *SPC*^Cre/+^;*Csf2*^fl^ and *Csf2*^Δ/Δ^ mice. (**B**) Flow cytometry analysis of CD45^−^tdTomato^+^ populations in P10 lungs of *Csf2*^fl^ and *SPC*^Cre/+^;*Csf2*^fl^ mice, gated on live cells. (**C**) Flow cytometry analysis of CD45^+^tdTomato^+^ populations in P10 lungs of *Csf2*^fl^ and *SPC*^Cre/+^;*Csf2*^fl^ mice, gated on live cells. (**D**) Total GM-CSF quantification in lung conditioned media from P10 *Csf2*^fl^, Vav1^iCre/+^;*Csf2*^fl^, *SPC*^Cre/+^;*Csf2*^fl^ and *Csf2*^Δ/Δ^ mice. (**E**) Flow cytometry analysis of AMs (CD11c^+^SiglecF^+^) in P10 lungs of *Csf2*^fl^ and *SPC*^Cre/+^;*Csf2*^fl^ mice, gated on live CD45^+^ cells. (**F**) AM (CD45^+^Ly-6G^−^SiglecF^+^CD11c^+^CD64^+^) quantification in P10 lungs of *Csf2*^fl^ and *SPC*^Cre/+^;*Csf2*^fl^ mice. (**A**) Data are from one experiment representative of two independent experiments. (**B+C, E)** Data pooled from 4 independent experiments. (**D**) Data from one independent experiment.

### Timed induction of *Csf2* expression in nascent AT2s initiates AM differentiation in fetal lung monocytes

Given the initiation of AM differentiation during fetal development (Guilliams et al., 2013; Schneider et al., 2014b), we speculated that the induction of GM-CSF expression by relevant sources may be timed with the appearance of AM precursors and the initiation of their local differentiation towards an AM fate. To establish a timeline of *Csf2* expression, we performed a kinetic analysis of *Csf2*-tdTomato reporter expression in perinatal mouse lungs. *Csf2*^+/+^ and *Csf2*^+/fl^ embryos and neonates were analyzed at E14.5, E16.5, E18.5, P0 and P4 to evaluate tdTomato signal in the hematopoietic and epithelial compartments. Few CD45^+^Lin^low/–^ tdTomato^+^ cells were detected during embryonic development, though there was a substantial increase following birth **(Supp. Fig. 4A**). This is consistent with the expansion of lung ILC2s during that time period (de Kleer et al., 2016; Saluzzo et al., 2017; Schneider et al., 2019; Steer et al., 2017). From E14.5 to P4, the vast majority of the tdTomato signal was detected in the CD45^−^EpCAM^+^ epithelial compartment. The first tdTomato^+^ cells were detected at E18.5, followed by a steady increase in the expression level and percentage of *Csf2*-reporter^+^ epithelial cells after birth (**Fig. 5A, B**). These results imply that epithelial *Csf2* expression is first induced in the lung between E16.5 and E18.5, concomitant with the commencement of AT2 differentiation (Treutlein et al., 2014), where it is then maintained constitutively.

**Figure 5.**
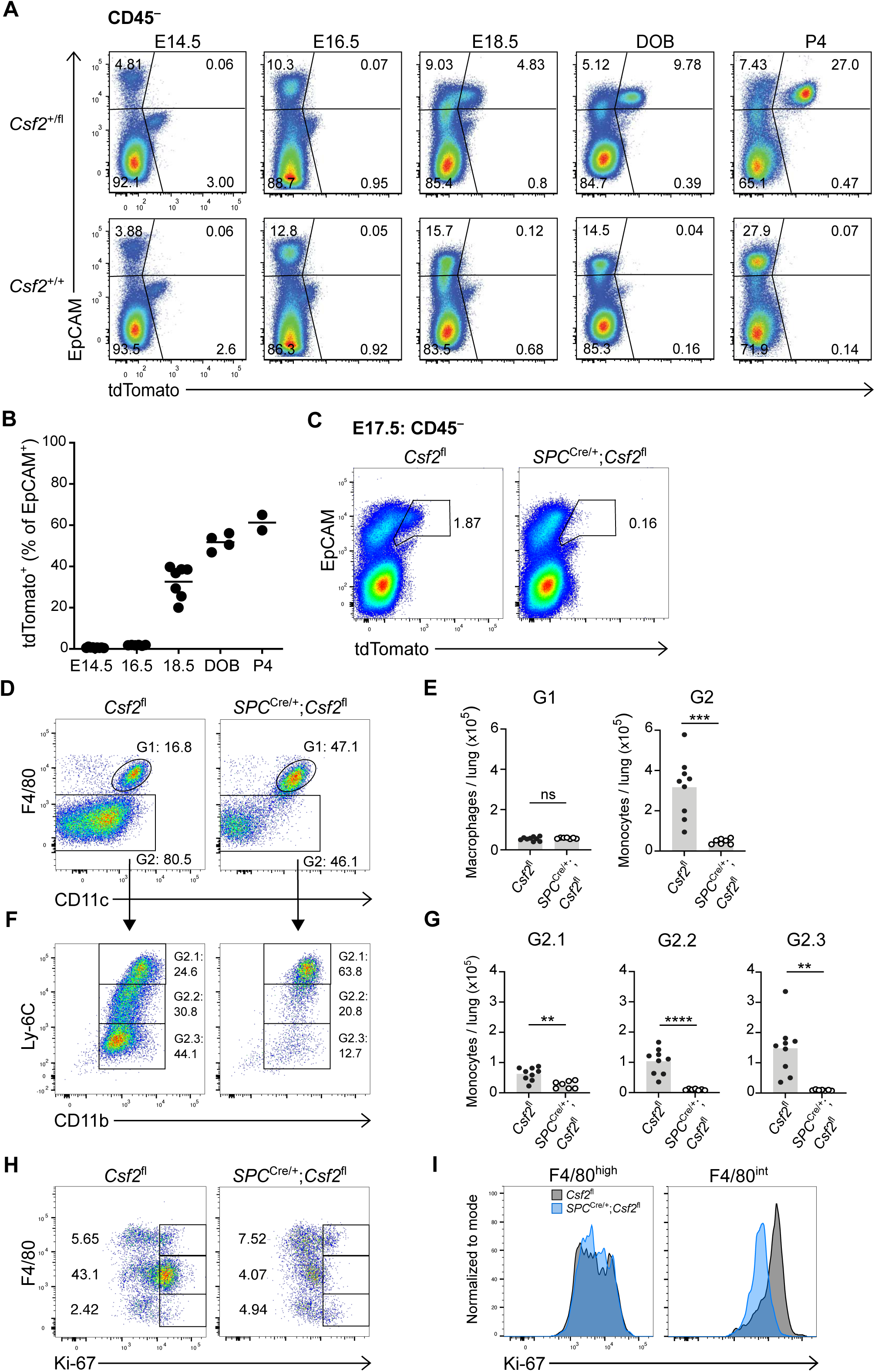
Timed induction of *Csf2* expression in nascent AT2s instructs AM differentiation in fetal lung monocytes. **(A)** Kinetics of tdTomato expression in the CD45^−^ compartment of perinatal lungs of *Csf2*^+/fl^ and *Csf2*^+/+^ mice ranging from E14.5 to P4, determined by flow cytometry. (**B**) Percentage of CD45^−^ EpCAM^+^tdTomato^+^ cells of mice in A. (**C**) Flow cytometry analysis of CD45^−^ tdTomato^+^ populations in E17.5 lungs of *Csf2*^fl^ and *SPC*^Cre/+^;*Csf2*^fl^ mice. (**D**) Flow cytometry analysis of primitive macrophages (G1; CD45^+^Ly-6G^−^CD64^+^MHCII^−^F4/80^high^) and fetal monocytes (G2; CD45^+^Ly-6G^−^CD64^+^MHCII^−^F4/80^int^) at E17.5 in the lungs of *Csf2*^fl^ and *SPC*^Cre/+^;*Csf2*^fl^ mice. (**E**) Quantification of primitive macrophages (G1) and fetal monocytes (G2) at E17.5 in the lungs of *Csf2*^fl^ and *SPC*^Cre/+^;*Csf2*^fl^ mice. (**F**) Further flow cytometry analysis of Ly-6C and CD11b levels in the developing fetal monocytes population (G2) in E17.5 lungs of *Csf2*^fl^ and *SPC*^Cre/+^;*Csf2*^fl^ mice. (**G**) Quantification of Ly-6C^high^ (G2.1), Ly-6C^int^ (G2.2), and Ly-6C^low^ (G2.3) fetal monocytes in E17.5 lungs of *Csf2*^fl^ and *SPC*^Cre/+^;*Csf2*^fl^ mice. (**H**) Flow cytometry analysis of Ki-67 expression in the CD45^+^ compartment of fixed E17.5 lungs in *Csf2*^fl^ and *SPC*^Cre/+^;*Csf2*^fl^ mice. (**I**) Further flow cytometry analysis of Ki67 expression levels in primitive macrophages (CD45^+^F4/80^high^) and fetal monocytes (CD45^+^F4/80^int^) at E17.5 in lungs of *Csf2*^fl^ (grey) and *SPC*^Cre/+^;*Csf2*^fl^ (blue) mice. (**A+B**) Data representative of at least two independent experiments per time point. (**B**) Data are pooled from multiple independent experiments. (**C, D, F**) Data are from one experiment, representative of three independent experiments. (**E, G**) Data are pooled from three independent experiments. (**H+I**) Data are from one experiment, representative of two independent experiments.

Previous reports identified the same time period as the beginning of fetal monocyte differentiation towards an AM identity (Guilliams et al., 2013; Schneider et al., 2014b). To assess if AT2-derived GM-CSF locally initiates these first steps of AM differentiation, we characterized the myeloid compartment in embryonic E17.5 lungs from *SPC*^Cre^;*Csf2*^fl^ mice. E17.5 was specifically chosen in order to capture the time point when *Csf2*-reporter expression first begins to emerge from CD45^−^EpCAM^+^ cells (**Fig. 5C**). Specific Cre activity was validated by the abrogation of this tdTomato expression (**Fig. 5C**), suggesting efficient deletion of *Csf2* from the AT2 lineage. Our results were consistent with prior studies reporting activity of the 3.7-kb human SP-C promoter in alveolar epithelial progenitor cells of the primordial lung buds (Wert et al., 1993). The minor *Csf2*-reporter expression among CD45^+^ cells was unchanged in the E17.5 lungs of *SPC*^Cre^;*Csf2*^fl^ mice (**Supp. Fig. 4B**). The macrophage/monocyte populations were then evaluated based on the canonical markers F4/80, CD11c, CD11b, and Ly-6C (**Supp. Fig. 4C)**. Primitive macrophages were identified as a relatively homogeneous population of F4/80^high^ cells that were equally abundant in *SPC*^Cre^;*Csf2*^fl^ and *Csf2*^fl^ embryos (**Fig. 5D, E**). In contrast, the F4/80^int^ cells comprise a heterogeneous compartment consisting of fetal monocytes that adopt a waterfall-shaped distribution of Ly-6C vs. CD11b, CD11c, and CD64 on their differentiation trajectory towards pre-AMs (**Fig. 5F, Supp. Fig. 4D, E**) (Guilliams et al., 2013; Schneider et al., 2014b). Notably, there was a significant reduction in these cells in *SPC*^Cre^;*Csf2*^fl^ mice predominantly affecting the Ly-6C^int^ and Ly-6C^low^ populations, which were almost completely absent (**Fig. 5E-G; Supp. Fig. 4D, E**). Further assessment of these early F4/80^int^ monocyte and F4/80^high^ macrophage populations for Ki-67 expression showed a significant and specific reduction in proliferation of fetal monocytes but not primitive macrophages in the *SPC*^Cre^;*Csf2*^fl^ lungs relative to their littermate controls (**Fig. 5H, I**).

Together, our results demonstrate that the intimate relationship between epithelial cells and AMs is already present in the fetal lung, where timed expression of GM-CSF from developing AT2s is the essential local signal to initiate robust *in situ* expansion of fetal monocytes and concomitantly, the first steps of AM differentiation.

### GM-CSF ablation in AT2s (but not hematopoietic cells) results in the absence of AMs and the development of PAP

Having identified AT2s as the critical source of GM-CSF for perinatal AM differentiation, we next asked whether this relationship perpetuates throughout adulthood. Given that diverse populations of hematopoietic cells in the lungs are capable of expressing GM-CSF, it is conceivable that their role in AM maintenance is only apparent following AT2-dependent AM development. Many immune cell subsets accumulate over the course of the first few weeks of life and are more numerous in the lungs of adult mice compared to the developing lungs of the early postnatal period (de Kleer et al., 2016). This influx of immune cells may remodel postnatal lung niches, affect the relative contribution of GM-CSF from individual sources, or possibly compensate for niche signals that may be missing. To determine whether the accumulation of immune cells in mature lungs modifies the AM-GM-CSF biology observed in neonates and embryos, we profiled myeloid populations in lungs of adult *Vav1*^iCre^;*Csf2*^fl^, *SPC*^Cre^;*Csf2*^fl^, *Csf2*^Δ/Δ^ and control mice. Evaluation of tdTomato expression by flow cytometry demonstrated hematopoietic-specific deletion of tdTomato signal in the presence of *Vav1*^iCre^, epithelial-specific deletion of tdTomato signal in the presence of *SPC*^Cre^, and loss of all tdTomato signal in the case of *Csf2*^Δ/Δ^ mice (**Fig. 6A**). Analysis of the adult myeloid compartment across the different genotypes duplicated the results found in neonatal lungs. As observed in the lungs of control mice, the AMs in *Vav1*^iCre^;*Csf2*^fl^ lungs were present at normal numbers, contrasting strongly with the lungs of both the *SPC*^Cre^;*Csf2*^fl^ and the *Csf2*^Δ/Δ^ mice, in which the AMs were completely absent (**Fig. 6B, C**). Notably, we again examined the effects of constitutive basophil-depletion on AM homeostasis. Similar to our observations in neonatal mice, basophils were absent from lungs of adult *Mcpt8*^YFP-Cre^;*R26*^DTA^ mice (**Supp. Fig. 5A, B**), whereas the number and surface marker profile of their AMs was indistinguishable from those of littermate control lungs (**Fig, 6D, Supp. Fig. 5C-F**). Protein and cholesterol concentrations in the bronchioalveolar lavage fluid (BALF) were also comparable between the two groups, suggesting functional AMs (**Fig 6E, F**). To further fortify the results observed with *Vav1*^iCre^ and *SPC*^Cre^-mediated *Csf2* deletion, we performed comprehensive spectral flow cytometry profiling and high-dimensional analysis of the myeloid compartments in these mice. The results showed normal representation of the AM cluster in control and *Vav1*^iCre^;*Csf2*^fl^ lungs, whereas this cluster in the lungs of both the *SPC*^Cre^;*Csf2*^fl^ and the *Csf2*^Δ/Δ^ mice was completely absent (**Fig. 6G**). Other myeloid populations could be identified based on their expression of signature markers and were largely unperturbed (**Fig. 6H**).

**Figure 6.**
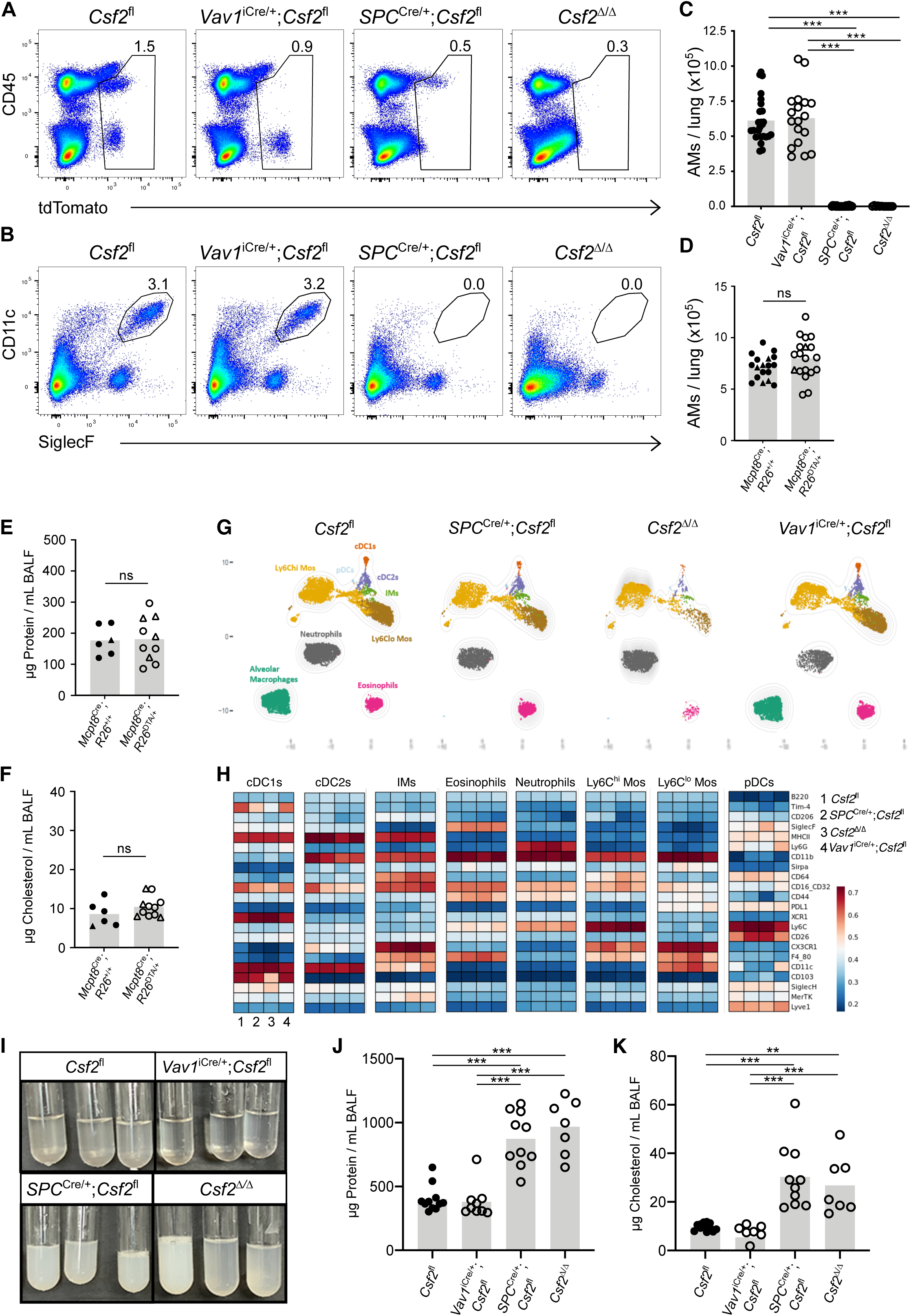
AT2-specific deletion of *Csf2* leads to AM depletion in adult lungs. (**A**) Flow cytometry analysis of tdTomato^+^ populations in adult *Csf2*^fl^, *Vav1*^iCre/+^;*Csf2*^fl^, *SPC*^Cre/+^;*Csf2*^fl^ and *Csf2*^Δ/Δ^ mice, gated on live cells. (**B**) Flow cytometry analysis of AMs (CD11c^+^SiglecF^+^) in adult *Csf2*^fl^, *Vav1*^iCre/+^;*Csf2*^fl^, *SPC*^Cre/+^;*Csf2*^fl^ and *Csf2*^Δ/Δ^ mice, gated on live CD45^+^ cells. (**C**) Quantification of AMs (CD45^+^Ly-6G^−^SiglecF^+^CD11c^+^CD64^+^) in adult lung of *Csf2*^fl^, *Vav1*^iCre/+^;*Csf2*^fl^, *SPC*^Cre/+^;*Csf2*^fl^ and *Csf2*^Δ/Δ^ mice. (**D**) AM (CD45^+^SiglecF^+^CD11c^+^) quantification in adult lungs of *Mcpt8*^YFP-Cre^;R26^+/+^ and *Mcpt8*^YFP-Cre^;R26^DTA/+^ mice. *Mcpt8*^YFP-Cre/+^ mice are indicated by circles, while *Mcpt8*^YFP-Cre/YFP-Cre^ mice are indicated by triangles. (**E**) Quantification of protein in BALF from adult *Mcpt8*^YFP-Cre^;R26^+/+^ and *Mcpt8*^YFP-Cre^;R26^DTA/+^ mice. Genotypes are indicated as per (D). (**F**) Quantification of total cholesterol in BALF from adult *Mcpt8*^YFP-Cre^;R26^+/+^ and *Mcpt8*^YFP-Cre^;R26^DTA/+^ mice. Genotypes are indicated as per (D). (**G**) A representative UMAP map showing the FlowSOM-guided meta-clustering of the myeloid compartment in *Csf2*^fl^, *SPC*^Cre/+^;*Csf2*^fl^, *Csf2*^Δ/Δ^ and *Vav1*^iCre/+^;*Csf2*^fl^ mice. (**H**) Heatmap displaying the median antigen intensity of markers used to generate (G). (**I**) BALF from adult *Csf2*^fl^, *Vav1*^iCre/+^;*Csf2*^fl^, *SPC*^Cre/+^;*Csf2*^fl^ and *Csf2*^Δ/Δ^ mice. (**J**) Quantification of protein in BALF from adult *Csf2*^fl^, *Vav1*^iCre/+^;*Csf2*^fl^, *SPC*^Cre/+^;*Csf2*^fl^ and *Csf2*^Δ/Δ^ mice. (**K**) Quantification of total cholesterol in BALF from adult *Csf2*^fl^, *Vav1*^iCre/+^;*Csf2*^fl^, *SPC*^Cre/+^;*Csf2*^fl^ and *Csf2*^Δ/Δ^ mice. (**A+B**) Data are from one experiment representative of four independent experiments. (**C**) Data are pooled from four independent experiments. (**D**) Data are pooled from three independent experiments. (**E+F**) Data are pooled from two independent experiments. **(G-I**) Data are from one experiment representative of two independent experiments. (**J+K**) Data are pooled from two independent experiments.

It has been well-documented in both humans and mice that missing or malfunctioning AMs, arising due to defects in GM-CSF signaling, results in the accumulation of cellular and surfactant debris within the lumen of the alveoli, a condition known as PAP (Trapnell et al., 2019). To further verify the importance of different GM-CSF sources for the homeostatic function of AMs, the BALF was collected from the lungs of *Vav1*^iCre^;*Csf2*^fl^ and *SPC*^Cre^;*Csf2*^fl^ mice, and analyzed with that from *Csf2*^Δ/Δ^ mice and pooled littermate control mice for signs of PAP. From both the flow cytometry plots and visual inspection of the extracted BALF, excess cellular debris was evident in the *SPC*^Cre^ and *Csf2*^Δ/Δ^ samples, but was not detectable in the *Vav1*^iCre^ nor the control samples (**Fig. 6I, and data not shown**). Quantification of total protein and cholesterol content of the BALF further indicated significantly higher levels of both protein and cholesterol in *SPC*^Cre^ and *Csf2*^Δ/Δ^ lungs relative to *Vav1*^iCre^ and control lungs (**Fig. 6J, K**). Together, these results suggest that loss of epithelial *Csf2* expression not only results in loss of the entire AM population, but also leads to PAP in the lungs of these mice. The presence of PAP further demonstrates that the lost AM population has not been functionally replaced in these AM-deficient lungs and it suggests that hematopoietic sources are neither able to compensate for the loss in GM-CSF production by AT2s, nor do they have a tangible effect on homeostatic AM function.

### Inducible AT2-specific ablation of *Csf2* expression in adult lungs results in AM population atrophy

Despite the clear reliance of developing AMs on AT2-derived GM-CSF, it was still not certain whether fully-developed AMs remained dependent on GM-CSF for their survival. Furthermore, constitutive absence of AMs and the resulting presence of PAP could impact *Csf2* expression by non-AT2s and thereby impede a definitive assessment of the critical GM-CSF source during adulthood. Our data thus demonstrated a requirement for assessing how mature AMs are maintained in the lungs. To address this query, we generated a conditional inducible mouse model in which *Csf2* expression from SP-C^+^ AT2 cells could be selectively depleted in adulthood upon treatment with tamoxifen. This was achieved by crossing *Csf2*^fl^ reporter mice with a tamoxifen-inducible *SPC*^CreERT2^ mouse line. *SPC*^CreERT2/CreERT2^;*Csf2*^fl^ mice were treated with tamoxifen, and lungs were analyzed by flow cytometry three weeks later (**Fig. 7A**). Evaluation of the tdTomato signal from epithelial CD45^−^EpCAM^+^ cells and hematopoietic CD45^+^Lin^−^Thy1.2^+^ cells revealed that there was specific decrease in the tdTomato signal from AT2s but no change in the hematopoietic-derived tdTomato signal in the presence of tamoxifen-activated *SPC*^CreERT2^ (**Fig. 7B, C**). This suggests that the tamoxifen treatment was specific for deletion of *Csf2* expression from the AT2s, and did not affect *Csf2* expression from the hematopoietic compartment. Three weeks post-tamoxifen treatment, AMs were found to be absent in the presence of activated *SPC*^CreERT2^ (**Fig. 7D, E**). Together these results suggest that even after the postnatal development of AMs is complete, AT2-derived GM-CSF continues to be a critical niche factor for the maintenance of AM in the adult alveoli. Furthermore, the crucial function of nourishing and maintaining AMs via GM-CSF production appears to be unique to the AT2 lineage.

**Figure 7.**
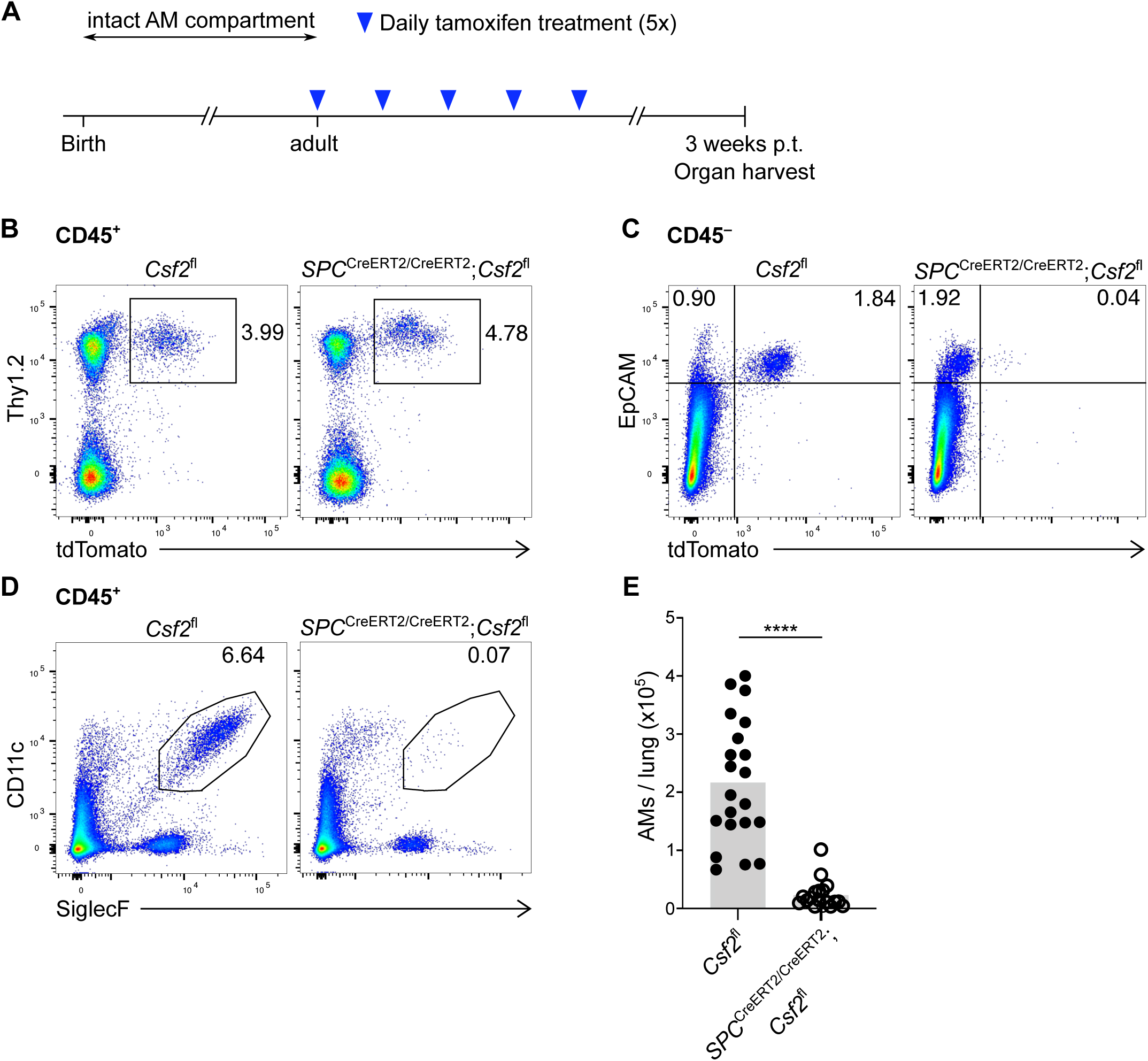
Inducible AT2-specific ablation of Csf2 expression in adult lungs results in AM population atrophy. Adult *Csf2*^fl^ and *SPC*^CreERT2/CreERT2^;*Csf2*^fl^ mice were treated with tamoxifen according to the timeline in (**A**). (**B, C**) Flow cytometry analysis of CD45^+^tdTomato^+^ (**B**) and CD45^−^tdTomato^+^ (**C**) populations in the lungs three weeks after tamoxifen administration, gated on live cells. (**D, E**) Flow cytometry analysis of AMs (CD11c^+^SiglecF^+^) gated on live CD45^+^ cells (**D**) and quantification of AMs (CD45^+^Ly-6G^−^ SiglecF^+^CD11c^+^CD64^+^) (**E**) three weeks after tamoxifen administration. (**B-D**) Data are from one experiment representative of three independent experiments. (**E**) Data are pooled from three experiments.

## DISCUSSION

Using a novel *Csf2* reporter allele and multiple conditional knockout strains, our study generated findings critical to the understanding of the AM niche in the developing and adult lung. First, we defined hematopoietic and non-hematopoietic sources of GM-CSF in the perinatal lung. Second, we demonstrated that the hematopoietic compartment, including ILC2s and basophils, are neither a necessary source of GM-CSF nor a critical compartment for AM development. Third, using multiparameter flow cytometry, we identified AT2s as the major, if not exclusive, non-hematopoietic pulmonary compartment with constitutive GM-CSF expression. Lastly, constitutive and inducible deletion of *Csf2* from the AT2 lineage revealed that AT2-derived GM-CSF plays a necessary and non-redundant function in promoting AM fate in fetal monocytes, in establishing the postnatal AM compartment, and in regulating AM maintenance in the adult lung.

Transgenic reporter alleles and scRNA-seq analysis have spurred major developments in the understanding of cytokine biology, enabling the identification of cytokine sources in tissues and guiding further *in vivo* validation. Here, we report the generation of a new GM-CSF reporter line suitable to investigating GM-CSF expression *in situ* and *ex vivo*. Furthermore, this *Csf2*^fl^ reporter can be readily converted to a conditional knockout allele by crossing the reporter mice to a Cre-expressing mouse line. The decrease in tdTomato signal after Cre-mediated recombination of the loxP-flanked *Csf2* exons, which we speculate occurs due to deletion of the associated splice acceptor sites and thus reduced expression of the truncated bicistronic transcript, enables a convenient readout of the recombination efficiency. Using this novel tool to study GM-CSF expression in the neonatal mouse lung, we identified multiple hematopoietic sources, including ILC2s, T cells and NK cells, among which ILC2s produced the most GM-CSF on a per-cell level. Our data is in agreement with prior reports, whose findings were based on *Csf2* fate-mapping and scRNA-seq analysis (Cohen et al., 2018; Komuczki et al., 2019)

Despite the significant homeostatic expression of GM-CSF by various pulmonary immune cells, our studies using *Vav1^i^*^Cre^-mediated *Csf2* deletion clearly demonstrate that hematopoietic-derived GM-CSF is dispensable for the postnatal differentiation and the subsequent maintenance of AMs. This finding was unexpected given the high *Csf2* expression by ILC2s and the reports suggesting that GM-CSF is included in the ILC2 effector-cytokine repertoire that is induced perinatally (Cohen et al., 2018; de Kleer et al., 2016; Saluzzo et al., 2017; Schneider et al., 2019), and which led to speculation about ILC2 functions in shaping homeostatic myeloid cell compartments beyond the regulation of eosinophils (Nussbaum et al., 2013). Furthermore, there are reports of important ILC-GM-CSF interactions in other contexts. Notably, ILC3-derived GM-CSF in the colon facilitates DC and macrophage-mediated T_reg_ differentiation and thereby contributes to oral tolerance (Mortha et al., 2014). Under conditions of tissue perturbation, ILC-mediated cross-talk via GM-CSF can amplify inflammation through activation of monocytes and eosinophils (Griseri et al., 2015; Song et al., 2015). Although not addressed by our studies, similar cellular interaction modules might also be active in the lung, including in allergic inflammation (Behrens et al., 2015; Nobs et al., 2021). The biological function of constitutive *Csf2* expression by pulmonary ILC2s, which we validated on an mRNA and protein level, however, remains unclear. It is possible that ILC2s create local tissue microenvironments that are enriched for GM-CSF and provide context-dependent support for particular myeloid compartments, such as eosinophils or DCs. Indeed, bone marrow ILC2s were recently reported to increase GM-CSF expression and to support hematopoietic recovery under conditions of stress hematopoiesis following treatment with 5-fluorouracil (Sudo et al., 2021). Further studies are thus required to reveal the more subtle consequences of lung hematopoietic *Csf2*-deficiency with the necessary spatial resolution.

The dispensability of hematopoietic GM-CSF for developing AMs was also surprising given a recent report that highlighted basophils as a critical regulator of AM development. In the developing lung, basophils were reported to express GM-CSF and to rely on GM-CSF receptor signaling; these cells were also suggested to critically regulate AM differentiation and numbers (Cohen et al., 2018). Our own investigations into basophil-AM biology using a genetic model of constitutive basophil depletion, however, did not confirm any changes in AM numbers nor any obvious defects in AM development, as determined by expression of signature surface markers, in the lungs of neonatal and adult mice. We can only speculate about the reason for these conflicting results. In addition to a genetic model of constitutive basophil depletion, Cohen *et al*. also employed an antibody-mediated basophil depletion strategy to functionally test the role of basophils *in vivo*. Newborn mice were treated intranasally with a relatively large amount of MAR-1 antibody, which was followed by lung cell analysis the following day. This may be a somewhat short time window to study basophil depletion-mediated consequences on AM development, though it does fall within the time window of fetal monocyte-to-AM differentiation. The MAR-1 antibody itself is known to bind and deplete FcεRIα-expressing cells including basophils, but concomitant antibody-mediated FcεRIα crosslinking has also been demonstrated to potently activate basophils, causing their degranulation and release of bioactive mediators *in vitro* and *in vivo* (Hübner et al., 2011; Pellefigues et al., 2019). Furthermore, a recent report demonstrated binding of MAR-1 to FcγRI and IV (Tang et al., 2019). Such off-target effects could alter gene expression in other cells, warranting caution when critically relying on the use of MAR-1 treatment. Although our studies using a genetic basophil-depletion strategy lack the high resolution needed to detect subtle transcriptional alterations in the AM compartment, our results cast doubt on the suggested importance of basophils for AM development.

Using comprehensive flow cytometry profiling, we identified AT2 cells as the major, if not exclusive, non-hematopoietic source of pulmonary GM-CSF under homeostatic conditions. Our embryonic analysis revealed that the production of epithelial GM-CSF, first detectable during late gestation and maintained across the entire lifespan, is acquired concomitantly with AT2 lineage commitment. The timepoint around which *Csf2* expression is first detected (E16.5-E18.5) corresponds to the transition from the canalicular stage of lung organogenesis (E15-E17) to the saccular stage of development (E17-P0) (Swarr and Morrisey, 2015). During this period, epithelial cells, which may derive from bipotent or pre-committed progenitors, begin to express transcriptional signatures reminiscent of mature AT1s and/or mature AT2s while they progressively differentiate towards nascent AT1 and AT2 lineages (Desai et al., 2014; Frank et al., 2019; Gonzalez et al., 2019; Treutlein et al., 2014; Zepp et al., 2021). Although our *Csf2* reporter is first detectable by flow cytometry in E17.5-E18.5 lungs, assigning the onset of fetal *Csf2* expression to a precise developmental stage is complicated by not only the dynamic epithelial compartment differentiation and the co-expression of AT1 and AT2 markers by progenitors (including SP-C), but also the differences in maturation kinetics between GM-CSF and tdTomato, the latter being a slow-maturing fluorescent protein (Balleza et al., 2018; Gehart et al., 2019). Further studies are needed to definitively identify the first *Csf2*-expressing epithelial cells in the fetal lung, and to determine whether these early GM-CSF producers are already committed to the AT2 lineage.

Our results also raise interesting questions about the regulation of this cytokine, which is constitutively expressed across vastly different cell types including AT2s, ILC2s and GM-CSF-producing T cells. Regulation of *Csf2* expression in epithelial cells is particularly intriguing, as it may be suggestive of tissue-immune crosstalk. Recent studies reported increased *Csf2* expression in alveolar and bronchial epithelial cells downstream of IL-1R signaling following *L. pneumophila* infection and house dust mite (HDM) treatment (Liu et al., 2020; Willart et al., 2012). Furthermore, reduced NF-κB signaling in alveolar epithelial cell cultures was found to be associated with their decreased *Csf2* expression (Sturrock et al., 2018). It will be interesting to explore whether similar pathways are involved in the onset of *Csf2* expression in early AT2s, and to investigate whether homeostatic regulation of epithelial GM-CSF production is altered after birth.

Unexpectedly given prior reports, we found that AMs are critically reliant not on hematopoietic GM-CSF but on epithelial GM-CSF derived from AT2s. Using genetic models of constitutive *SPC*^Cre^-mediated and inducible *SPC*^CreERT2^-mediated deletion of *Csf2*, lungs from fetal, neonatal and adult timepoints demonstrate that regulation of epithelial GM-CSF-AM-AT2 biology occurs on multiple levels. GM-CSF from early *SPC*^Cre^-expressing epithelial cells or their progeny (likely bipotent epithelial progenitors or nascent AT2s) supports the proliferation and local expansion of fetal monocytes. These c-Myb-dependent myeloid progenitors can first be detected in lungs around E13 (Hoeffel et al., 2015; Schulz et al., 2012), several days before the first signs of GM-CSF expression based on our kinetic analysis. The mechanism of this anticipatory positioning of AM progenitors is unclear and our findings suggest that additional pathways may exist to ensure their survival until lung epithelial-derived GM-CSF drives local expansion and instructs differentiation towards an AM fate. We found that this process is blocked in *SPC*^Cre/+^;*Csf2*^fl^ mice, which replicates the phenotype previously reported for global *Csf2*^−/−^ mice (Schneider et al., 2014b), and that completion of AM differentiation postnatally is solely dependent on GM-CSF from nascent and mature AT2s. Thus, our studies provide further evidence for the existence of discrete tissue niches that critically regulate cell fate of developing tissue-resident immune cells (Guilliams and Scott, 2017). Additionally, constitutive GM-CSF production from AT2s is required for the maintenance of a mature AM compartment; our results reveal that AMs disappear within a few weeks of ablating AT2-specific *Csf2* expression in adult mice. Indeed, a role for GM-CSF in regulating AM self-renewal was suggested based on the capacity to stimulate the proliferation of isolated AM *in vitro* (Chen et al., 1988). It remains to be determined if this loss of AMs occurs due to abrogated self-renewal or enhanced cell death, the latter possibly caused by decreased expression of anti-apoptotic factors as suggested by studies using isolated AMs (Draijer et al., 2019).

Our studies unequivocally demonstrate the strict requirement for AT2-derived GM-CSF in AM development and maintenance. They also reveal the limitations of inferring ligand-receptor pairs and interaction modules primarily based on transcriptional data from dissociated organs lacking spatial context. However, emerging high-resolution spatially resolved transcriptomics, which enables the mapping of gene expression data to protein and structural information, will likely prove very useful for predicting cellular interactions in complex tissues with better precision via an unbiased systems-level approach (Marx, 2021). In the lung, the demand for a niche factor produced by a discrete source, despite its even higher expression in other cell types that fail to compensate, is remarkable. For GM-CSF and AMs, this may in part be explained simply by the higher abundance of AT2s compared to ILC2s. The total number of AT2s in an adult mouse lung is estimated to be approximately 10^7^ (Dzhuraev et al., 2019) while that of ILC2s may be less than 10^5^ (Ricardo-Gonzalez et al., 2018). In addition, AT2s are ideally positioned for intimate interaction with AMs, and organization of the AT2 secretion machinery might favor apical release of GM-CSF directly into the alveolar lumen where AMs reside. In contrast, ILC2s are found in collagen-rich adventitial niches that localize to bronchovascular areas, and they are rarely found in the alveolar parenchyma (Dahlgren et al., 2019). Therefore, ILC2-derived GM-CSF may be spatially segregated from AMs and consumed locally by other cell types. The processes that drive such niche organization and positioning of differentiating tissue-resident immune cells during organogenesis are largely unknown. It is intriguing to speculate that circulating fetal monocytes are recruited to the luminal side of the saccule by gradients of GM-CSF, where the highest GM-CSF concentration exists in what will later become the luminal side of mature alveoli and the niche environment for AMs.

Collectively, our study unveils AT2s as a constitutive and dominant source of GM-CSF in the alveolar niche, nurturing AMs in prenatal as well as postnatal stages of lung development. This AT2-GM-CSF-AM relationship persists from last gestation throughout adulthood, such that consistent AT2-derived GM-CSF is crucial for both AM development and maintenance.

## AUTHOR CONTRIBUTIONS

J.G. and S.S. conceived the study, designed and performed experiments, analyzed data and wrote the manuscript with C.S. F.R. performed additional experiments and assisted with data analysis. X.F. helped with experiments. H.-E.L. designed the original tdTomato reporter cassette, performed the ES cell work for *Csf2*^fl^ mice, and provided scientific input. R.M.L. acquired funding and provided resources for *Csf2* targeting, and provided scientific input. B.B. assisted with data analysis and provided scientific input. C.S. generated *Csf2*^fl^ mice together with H.-E.L., directed the study and wrote the paper with J.G. and S.S.

## ACKNOWLEDGMENTS

We thank Brigitte Herzog for technical expertise; M. Kopf for helpful discussion and support with mouse rederivation; J. Rock and H. Chapman for providing mice; C. Wagner for providing the anti-NaPi-IIb; and members of the Schneider laboratory for helpful discussions. This work was supported by grants from the UCSF Diabetes Research Center, US National Institutes of Health (NIH), the Howard Hughes Medical Institute, and the Sandler Asthma Basic Research (SABRe) Center at the UCSF to R.M.L, the Swiss National Science Foundation (310030_188450) to B.B., and the Peter Hans Hofschneider Professorship for Molecular Medicine to C.S.

## MATERIALS AND METHODS

### CONTACT FOR REAGENT AND RESOURCE SHARING

Further information and requests for resources and reagents should be directed to and will be fulfilled by the Lead Contact, Christoph Schneider.

#### Mice

*S*ftpcCre (*SPC*^Cre^) (Tg(Sftpc-Cre)1Blh) (Okubo and Hogan, 2004) mice were generously provided by J. Rock via R. Locksley. *S*ftpcCreERT2 (*SPC*^CreERT2^) knock-in mice (Sftpctm1(Cre/ERT2,rtTA)Hap (Chapman et al., 2011) were generously provided by H. Chapman via R. Locksley. *Vav1*^iCre^ (B6.Cg-Commd10Tg(Vav1-iCre)A2Kio/J; 008610,) mice were purchased from the Jackson Laboratory. *Mcpt8*^YFP-Cre^;*R26*^DTA^ mice were previously described (Sullivan et al., 2011) and provided by R. Locksley.

*Csf2*^fl^ mice were generated by homologous gene targeting in C57BL/6 embryonic stem cells using a previously described targeting strategy and reporter cassette that was modified to remove the BGHpA (PMID 26675736). Briefly, a pKO915-DT (Lexicon Genetics)-based targeting vector was generated with the following components encoded downstream of the DTA-SV40pA: a 2.6 kb 5’ homology arm, a loxP site and the complete third and fourth exon of *Csf2*, a reporter cassette, and a 3.2 kb 3’ homology arm beginning in the 3’ UTR of *Csf2*. The reporter cassette encoded (in order from 5’ to 3’) a loxP site, encephalomyocarditis virus internal ribosome entry site (IRES), tdTomato, and an frt-flanked neomycin resistance cassette. The final construct was linearized with NotI and transfected by electroporation into C57BL/6 embryonic stem cells. Cells were grown on irradiated feeders with the aminoglycoside G418 in the media, and neomycin-resistant clones were screened for 5’ and 3’ homologous recombination by PCR. Positive clones were selected and screened by PCR using the Expand Long Range dNTPack (Sigma), PCR buffer supplemented with 2% DMSO and the primer pairs for the 5’ region (LR 5’ fw: 5’-AATCCAGTTCACGCAGCTTGTACG-3’; LR 5’ rev: 5’-CCTTCAGCCCCTTGTTGAATACGC-3’) and the 3’ region (LR 3’ fw: 5’-TCGCCTTCTATCGCCTTCTTGACG-3’; LR 3’ rev: 5’-GATTTCCATCCATCTGAGACGGTG-3’) of the modified locus. Clones were injected into albino C57BL/6 blastocysts to generate chimaeras, and the male pups with highest ratios of black-to-white coat color were selected to breed with homozygous Gt(ROSA26)FLP1/FLP1 females (Jackson Laboratories; 009086) to excise the neomycin resistance cassette. Deletion of neomycin was confirmed by PCR using fw 5’-GGCTACTACTACGTGGACAC-3’ and rev 5’-CCAAGTTCCTGGCTCATTAC-3’. Offspring with germline transmitted and neo-deleted *Csf2*^fl^ allele were backcrossed to C57BL/6 mice to cross out the FLP1 allele. *Csf2*^fl^ genotyping was done using primers loxP fw: 5’-GCTTTTGAAATAGTGCTTCCCCAC-3’; loxP rev: 5’-AGGTTCCCAGTTCCAAGTGCTGTC-3’ (163 bp wild-type band; 131 bp floxed band). Deletion of the floxed allele was detected by PCR using primers “loxP fw” and “LR 5’ rev”, yielding 526 bp recombined or 1811 bp floxed products. *Csf2*^Δ/Δ^ mice with global *Csf2* deletion were generated from *SPC*^Cre/+^;*Csf2*^fl^ mice, in which Cre is active with high frequency in the male germline resulting in transmission of a recombined allele. Because of repeated germline Cre activity in both male and female *Vav1*^iCre^ mice, it is possible to have transmission of a *Csf2*^Δ^ allele to offspring. The germline-deleted allele, however, could not be readily distinguished from the *Vav1*^iCre^ recombined allele in biopsy samples due to the presence of hematopoietic cells in the biopsies. Thus all conditional KO groups were compared to their littermate Cre^−^ controls, and included both *Csf2*^fl/fl^ and *Csf2*^Δ/fl^ mice. No obvious differences in AM were observed between *Csf2*^Δ/fl^ and *Csf2*^fl^ mice (data not shown). All mice were generated on or previously backcrossed to the C57BL/6 background. Mice were bred and housed at the University of Zurich Laboratory Animal Sciences Center (LASC) Zurich, Switzerland, in an SOPF barrier facility. Experimental animals were internally transferred to an SPF barrier facility and were housed in individually ventilated cage units. Animal experiments were reviewed and approved by the cantonal veterinary office of Zurich (permit numbers ZH134/2019, ZH117/2019). Experiments were carried out using age- and sex-matched groups whenever possible.

#### Mouse treatments

Adult *SPC*^CreERT2^;*Csf2*^fl^ mice were treated with tamoxifen dissolved at 20 mg/mL in corn oil (Sigma). 20 μg tamoxifen per gram body weight were administered by oral gavage once a day over the course of five days. Mice were then sacrificed three weeks after the last tamoxifen treatment.

#### Tissue processing for flow cytometry

Unless indicated otherwise, all adult mice were euthanized by lethal intraperitoneal (i.p.) pentobarbital injection; pups were euthanized by either live decapitation or lethal i.p. pentobarbital injection. Lungs were perfused through the right cardiac ventricle with cold PBS and incubated in ice-cold RMPI-1640 until time of dissociation. Lungs underwent an initial mechanical dissociation using the m_lung_01_01 program on the gentleMACS dissociator (Miltenyi Biotec). Lung tissues were then digested with 50 μg/mL Liberase TM (Roche) and 25 μg/mL DNase I (Roche) in pre-warmed RPMI-1640 for 35 minutes at 37 °C with gentle rocking. This was followed by further dissociation using the m_lung_01_02 program on the gentleMACS dissociator. Single-cell suspensions were obtained after passing the homogenized samples through 70 μm cell strainers. Following red blood cell lysis (10x BD Pharmlyse), cells were washed, filtered, and stained for flow cytometry.

Embryonic lungs were harvested from timed-pregnant dams at various timepoints ranging from E14.5 to E18.5. After euthanization of the dam by CO_2_ inhalation, embryos were removed and placed in ice-cold PBS until time of dissection. For E16.5-E18.5 embryos, lungs were perfused intracardially with cold PBS prior to removal. In all cases, dissected lungs were incubated in cold FACS buffer until time of dissociation. Tissue was manually dissociated with scissors, and digested with 50 μg/mL Liberase TM (Roche) and 25 μg/mL DNase I (Roche) in pre-warmed RPMI-1640 for 15-20 minutes at 37°C with gentle rocking. Partially digested tissue was triturated with a P1000 pipette, and further digested for 15-20 minutes at 37 °C with gentle rocking. Upon completion of the digestion, tissue was triturated and filtered through 50 μm cell strainers. Following red blood cell lysis (10x BD Pharmlyse), cells were washed, filtered, and stained for flow cytometry.

#### Flow cytometry

Fc block (anti-mouse CD16/32, 2.4G2) was purchased from BioXCell. Anti-mouse CD45 (30-F11), and rat anti-mouse SiglecF (E50-2440) were purchased from BD Bioscience; anti-mouse I-A/I-E (M5/114.15.2), anti-mouse CD31 (390), anti-mouse CD104 (346-11A), anti-mouse CD140a (APA5), anti-mouse CD326 (G8.8), anti-mouse CD49f (GoH3), anti-mouse Podoplanin (8.1.1), anti-mouse Ly-6A (D7), anti-mouse CD11c (N418), anti-mouse XCR1 (ZET), anti-mouse CD11c (M1/70), anti-mouse CD103 (2E7), anti-mouse Ly-6G (1A8), anti-mouse Ly-6C (HK1.4), anti-mouse CD64 (X54-5/7.1), anti-mouse F4/80 (BM8), anti-mouse GR-1 (RB6-8C5), anti-mouse NK1.1 (PK136), anti-mouse CD8a (53-6.7), anti-mouse CD196 (29-2L17), anti-mouse CD19 (6D5), anti-mouse CD4 (RM4-5) anti-mouse CD90.2 (30-H12), anti-mouse FcER1a (MAR-1), anti-mouse CD3 (17A2), anti-mouse TCRb (H57-597), anti-mouse CD49b (DX5), anti-mouse TCRgd (GL3), and anti-mouse SIRPɑ (P84) were purchased from BioLegend; anti-mouse NKp46 (29A1.4) was purchased from Invitrogen; anti-mouse ST2 (DJ8) was purchased from MD Bioproducts. Lineage (Lin)^-^ cells were defined as lacking CD11b, CD11c, NK1.1, F4/80, Gr-1, FcεRIα, and Ter119. Dead cells were identified using DAPI, fixable violet Live/Dead (Thermo) or ZombieRed (Biolegend). For intracellular staining of proteins, cells were fixed using either 4% PFA (Electron Microscopy Science), or in the case of Ki-67 staining, the FoxP3 fixation kit. Following fixation, cells were washed and stained in permeabilization buffer (ThermoFisher, 00-8333-56) with either (1) anti-mouse Ki-67 (SolA15, ThermoFisher), or (2) primary anti-dsRed (Sicgen, AB8181-200, 1:200) together with primary rabbit anti-proSP-C (Sigma, AB3786, 1:1000) or primary rabbit anti-NaPi-IIb (provided by C. Wagner, (Hilfiker et al., 1998), 1:400), followed by a wash and secondary donkey anti-goat IgG-Alexa Fluor 555 (ThermoFisher, A-32816, 1:4000) and donkey anti-rabbit IgG-Alexa Fluor 488 (ThermoFisher, A-21206, 1:4000).

Samples were analyzed on a FACSymphony (BD Biosciences) with five lasers (355 nm, 405 nm, 488 nm, 561 nm, and 639 nm). For doublet exclusion, samples were gated by FSC-H and FSC-A, followed by SSC-H and SSC-A; FSC-A and SSC-A gating was used to exclude debris, followed by dead cells exclusion. Data analysis was performed using FlowJo (Treestar).

#### High-dimensional Analysis

For spectral cytometry data, samples were analyzed on a Cytek Aurora (Cytek Biosciences) with five lasers (355 nm, 405 nm, 488 nm, 561 nm, and 640 nm). The compensation matrix was corrected using FlowJo software (Tree Star). After live, single, CD45^+^ and lin^−^ (NK1.1^−^, CD19^−^, CD3^−^, Ter119^−^) and compensated cells were exported and imported into R Studio. Before automated high-dimensional data analysis, spectral cytometry data were transformed using an inverse hyperbolic sine (arcsinh) function with a cofactor in the range of between 1200 and 3500. Additionally, all spectral cytometry data were normalized between 0 and 1 to the 99.999th percentile of the merged sample.

#### Automated Population Identification

To identify myeloid immune cell populations accurately, we first carried out a step of FlowSOM clustering to generate a starting point of 100 nodes on pre-processed and combined flow cytometry datasets (Hartmann et al., 2016; Van Gassen et al., 2015). This was then followed by expert-guided manual meta clustering, using parameters listed in each figure legend. The respective k-value was manually chosen (in the range of between 7 and 12); identified clusters were annotated and merged based on a similarity of antigen expression in order to uphold the biological relevance of the dataset. Heatmaps display median expression levels of all markers per merged population and plotted using the R package ‘‘pheatmap’’ (Kolde, 2019). For data visualization, we used Uniform Manifold Approximation and Projection (UMAP) (McInnes et al., 2020). To create a UMAP of myeloid cells, we pooled equally proportioned 50,000 CD45^+^lin^−^ cells from each genotype.

#### Cell sorting

Major populations of cells expressing the GM-CSF reporter were sorted from homogenized lungs that were pooled from 2-3 individual *Csf2*^+/fl^ P10 mice. The three cell populations collected were ILC2s (tdTomato^+^CD45^+^Lin^−^CD3^−^Thy1.2^+^ST2^+^), T cells (tdTomato^+^CD45^+^CD3^+^), and epithelial cells (tdTomato^+^CD45^−^EpCAM^+^). Cells were sorted directly into RLT Lysis Buffer (Qiagen) using a FACSAria II (BD Bioscience). Each channel was loaded with 5’000-100’000 cells from each sample.

#### Quantitative RT–PCR

RNA was isolated from FACSorted cells using the Quick-RNA Microprep kit (Zymo) and reverse transcribed using the SuperScript IV VILO Master Mix with ezDNase Enzyme (ThermoFisher). The resulting cDNA was used as template for quantitative PCR (qPCR) with the Power SYBR Green reagent on a 7500 Fast Real-Time PCR System (Applied Biosystems). Transcripts were normalized to *Rps17* (40S ribosomal protein S17) expression, and relative expression was shown as 2^-ΔCt^. Primer sequences: *Rps17*, 5’-CGCCATTATCCCCAGCAAG-3’, 5’-TGTCGGGATCCACCTCAATG-3’; *Csf2* 5’-ACATGACAGCCAGCTACTAC-3’, 5’-TCAAAGGGGATATCAGTCAG-3’;

#### Bronchoalveolar Lavage (BAL)

Mice were euthanized by lethal i.p. pentobarbital injection. The trachea was exposed and cannulated with a 20G catheter (Surflo). 1 mL of cold PBS was instilled intratracheally; without removing the catheter or syringe, the lung was flushed three times with approximately 0.7 mL of the instilled PBS. Total recovered volume was approximately 0.8 mL/mouse. The BAL fluid (BALF) was then centrifuged at 1500 RPM for 5 minutes at 4 °C. The supernatant was collected and stored at -80 °C until further use. The cell pellets were washed with 2% FBS and 0.05% sodium azide in PBS and stored on ice until time of staining. Total protein in BALF was quantified using the Pierce BCA protein assay kit (ThermoFisher, 23225), as per manufacturer’s instructions. Total cholesterol in BALF was quantified using the Cholesterol/Cholesteryl Ester Assay Kit (Abcam, ab65359) as per manufacturer’s instructions.

#### Conditioned lung medium for cytokine detection

P10 pups were euthanized by lethal i.p. pentobarbital injection; lungs were then removed under aseptic conditions and incubated in a 24-well plate with 1.5 mL of complete RPMI media (RPMI-1640, 10% FBS, 1 mM HEPES, 100 U/mL/100 ug/mL Pen/Strep, 55 μM 2-mercaptoethanol) at 37 °C and 5% CO_2_. Lung conditioned media was harvested after 24 hours and stored at -80 °C until analysis. Cytokine concentration in lung conditioned media was measured using the Biolegend Legendplex Assay (740134). The assay was performed according to the manufacturer’s instruction, with the exception that 8 μL of sample, standard, assay buffer, beads, detection antibodies and Streptavidin-PE were used instead of the recommended 25 μL.

#### Immunofluorescence (IF)

Mice were euthanized by lethal i.p. pentobarbital and the trachea was exposed. Lungs were instilled intratracheally with 1 mL of ice cold 4% PFA followed by prewarmed 0.5% low-melting agarose. Lungs were covered with ice until the agarose solidified; removed; and incubated in cold 4% PFA for 3 hours at 4 °C in the dark. Samples were dehydrated in 30% sucrose for 24 hours at 4°C, embedded in OCT Cryo Media (Medite Inc.) on dry ice, and stored at -80 °C.

Frozen lungs embedded in OCT Cryo Media (Medite Inc.) were cut at the Cryostat (Leica CM3050) into 8 μm thick sections and placed on glass slides. Before staining, sections were washed for 5 minutes in PBS, circled with a hydrophobic PAP pen (Dako pen) and blocked with blocking buffer for one hour at room temperature. After blocking, samples were washed for 5 minutes in PBS and stained with primary antibodies rabbit anti-dsRed (1/250, Rockland, 600-401-379) and goat anti-CD45 (1/200, R&D System, AF114) for 1 hour at room temperature. Sections were washed 3 x 5 minutes in PBS and stained with secondary antibodies donkey anti-rabbit Alexa Fluor 555 (1/1000, Invitrogen, A-31572) and donkey anti-goat Alexa Fluor 488 (1/1000, Invitrogen, A-11055) for 1 hour at room temperature. Thereafter, sampled were washed 3 x 5 minutes in PBS and stained with DAPI for 10 minutes. After a last wash with PBS for 5 minutes, tissues were mounted using Vectashield Antifade Mounting Medium (Vecta Laboratories, Burlingame, California, United States). Fluorescence was detected with a Leica confocal microscope (Leica SP8 inverse; Leica Camera AG, Wetzlar, Germany). Leica LAS X software was used to acquire and process the images.

### QUANTIFICATION AND STATISTICAL ANALYSIS

All experiments were performed using randomly assigned mice without investigator blinding, with an aim of at least n=3 per group per biological replicate and at least 2 biological replicates per experiment (unless otherwise stated). No statistical methods were used to predetermine sample size. All data points and n values reflect biological replicates. No data were excluded, unless there was a major experimental error that justified the exclusion. Statistical analyses of two experimental groups were performed using unpaired, two-tailed Student’s t-tests; if the p-value was <0.05, results were deemed statistically significant. When statistical analyses of more than two experimental groups were required, ordinary one-way ANOVAs were employed; further analyses of statistically significant ANOVA results (p<0.05) were performed using a Tukey’s multiple comparisons test. In the case of two independent variables (condition and genotype), a two-way ANOVA was employed; further analyses of statistically significant ANOVA results (p<0.05) were performed using a Tukey’s multiple comparisons test. Intragroup variation was not assessed, except in the case of *Mcpt8*^YFP-Cre/+^ and *Mcpt8*^YFPCre/Cre^ mice, which allowed for these two genotypes to be pooled and experimental groups to be assigned solely on the status of the *Rosa26*^DTA^ allele. All statistical analysis was performed using Prism 8 (GraphPad Software).

---

**Figure S1.**
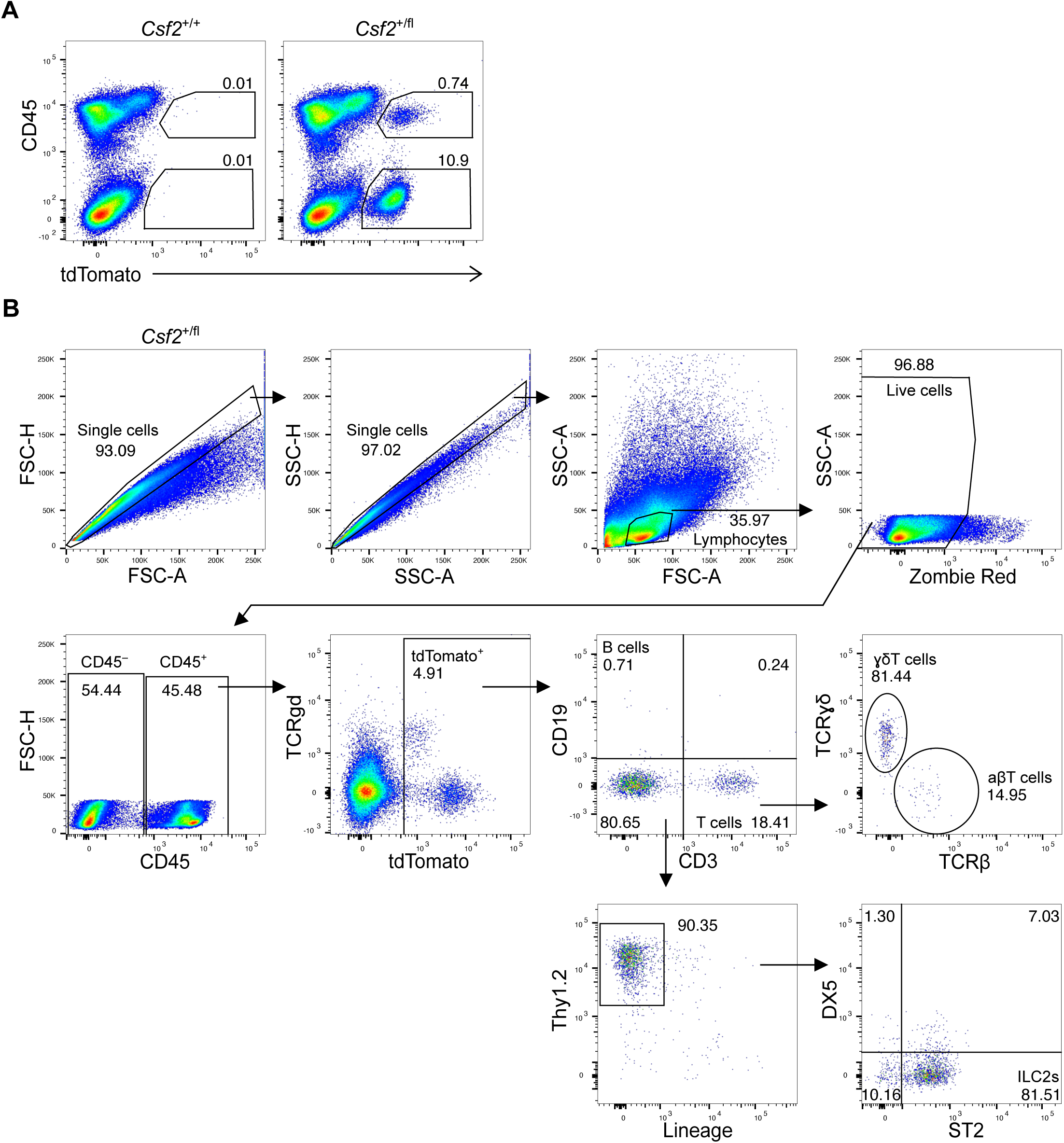
Relative GM-CSF contribution from the hematopoietic and non-hematopoietic compartments in the neonatal lung. (**A**) Flow cytometry of tdTomato expression by CD45^+^ and CD45^−^ cells in *Csf2*^+/+^ and *Csf2*^+/fl^ P10 lungs, gated on live cells. (**B**) Gating strategy used to identify lymphocytes in Fig. 1.

**Figure S2.**
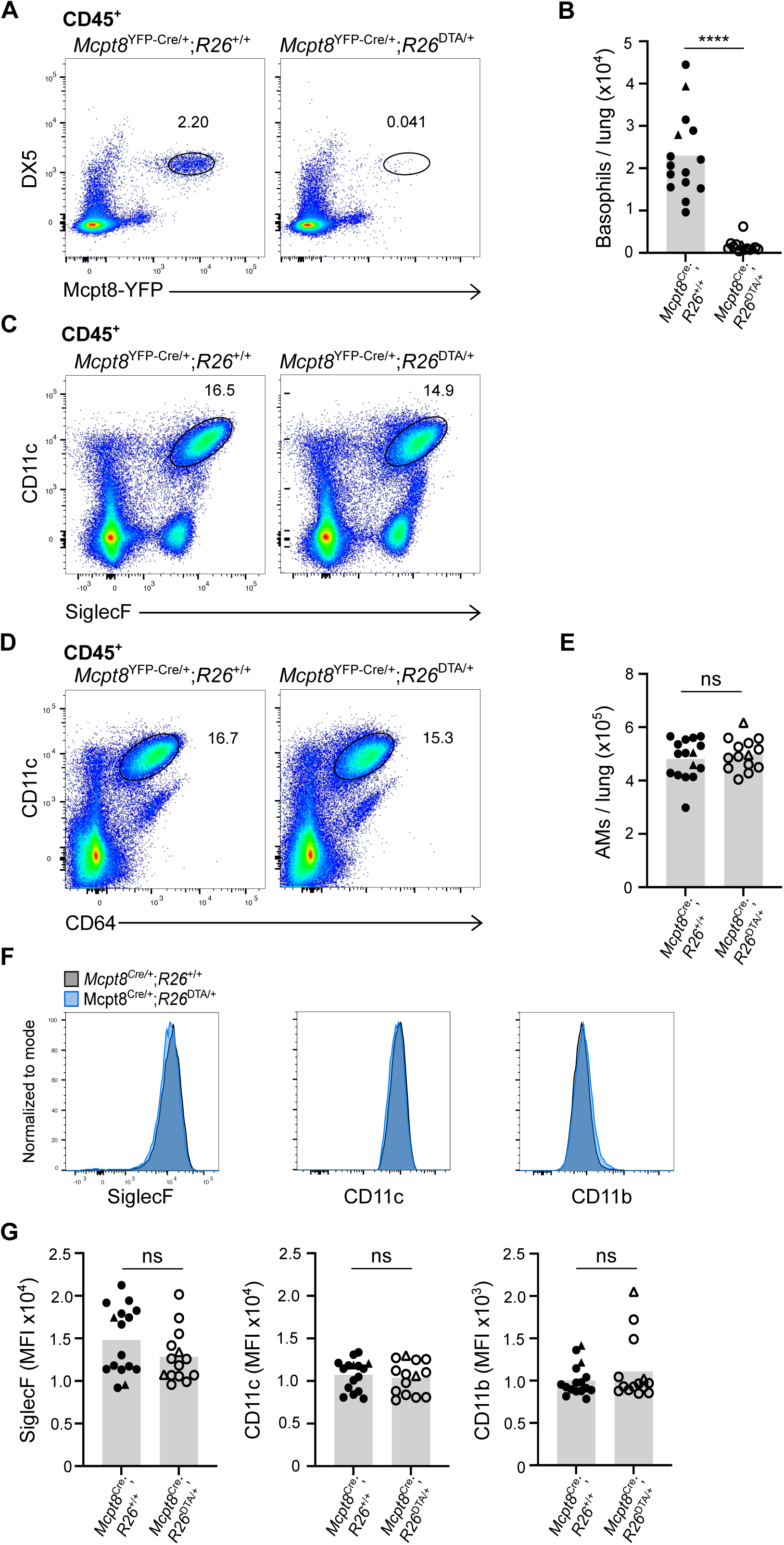
Constitutive depletion of basophils does not significantly alter the AM population in the neonatal lung. (**A**) Flow cytometry analysis CD45^+^DX5^+^Mcpt8-YFP^+^ basophils in P10 lungs of *Mcpt8*^YFP-Cre^;R26^+/+^ and *Mcpt8*^YFP-Cre^;*R26*^DTA/+^ mice. (**B**) Quantification of CD45^+^DX5^+^Mcpt8-YFP^+^ basophils in P10 lungs of *Mcpt8*^YFP-Cre^;*R26*^+/+^ and *Mcpt8*^YFP-Cre^;*R26*^DTA/+^ mice. *Mcpt8*^YFP-Cre/+^ mice are indicated by circles, while *Mcpt8*^YFP-Cre/YFP-Cre^ mice are indicated by triangles. (**C**) Flow cytometry analysis of AMs (CD45^+^CD11c^+^SiglecF^+^) in P10 lungs of *Mcpt8*^YFP-Cre^;*R26*^+/+^ and *Mcpt8*^YFP-Cre^;*R26*^DTA/+^ mice. (**D**) Flow cytometry analysis of AMs (CD45^+^CD11c^+^CD64^+^) in P10 lungs of *Mcpt8*^YFP-Cre^;*R26*^+/+^ and *Mcpt8*^YFP-Cre^;*R26*^DTA/+^ mice. (**E**) Quantification of AMs (CD45^+^CD11c^+^CD64^+^) in P10 lungs of *Mcpt8*^YFP-Cre^;*R26*^+/+^ and *Mcpt8*^YFP-Cre^;*R26*^DTA/+^ mice. Genotypes are indicated as per (B). (**F**) Expression levels of SiglecF, CD11c and CD11b for CD45^+^CD11c^+^CD64^+^ AMs at P10, as determined by flow cytometry, for *Mcpt8*^YFP-Cre^;*R26*^+/+^ (grey) and *Mcpt8*^YFP-Cre^;*R26*^DTA/+^ (blue) mice. (**G**) Quantification of SiglecF, CD11c and CD11b for AMs (CD45^+^CD11c^+^CD64^+^) at P10 for *Mcpt8*^YFP-Cre^;*R26*^+/+^ and *Mcpt8*^YFP-Cre^;*R26*^DTA/+^ mice. Genotypes are indicated as per (B). (**A, C, D, F**) Data are from one experiment, representative of three independent experiments. (**B, E, G**) Data are pooled from three independent experiments.

**Figure S3.**
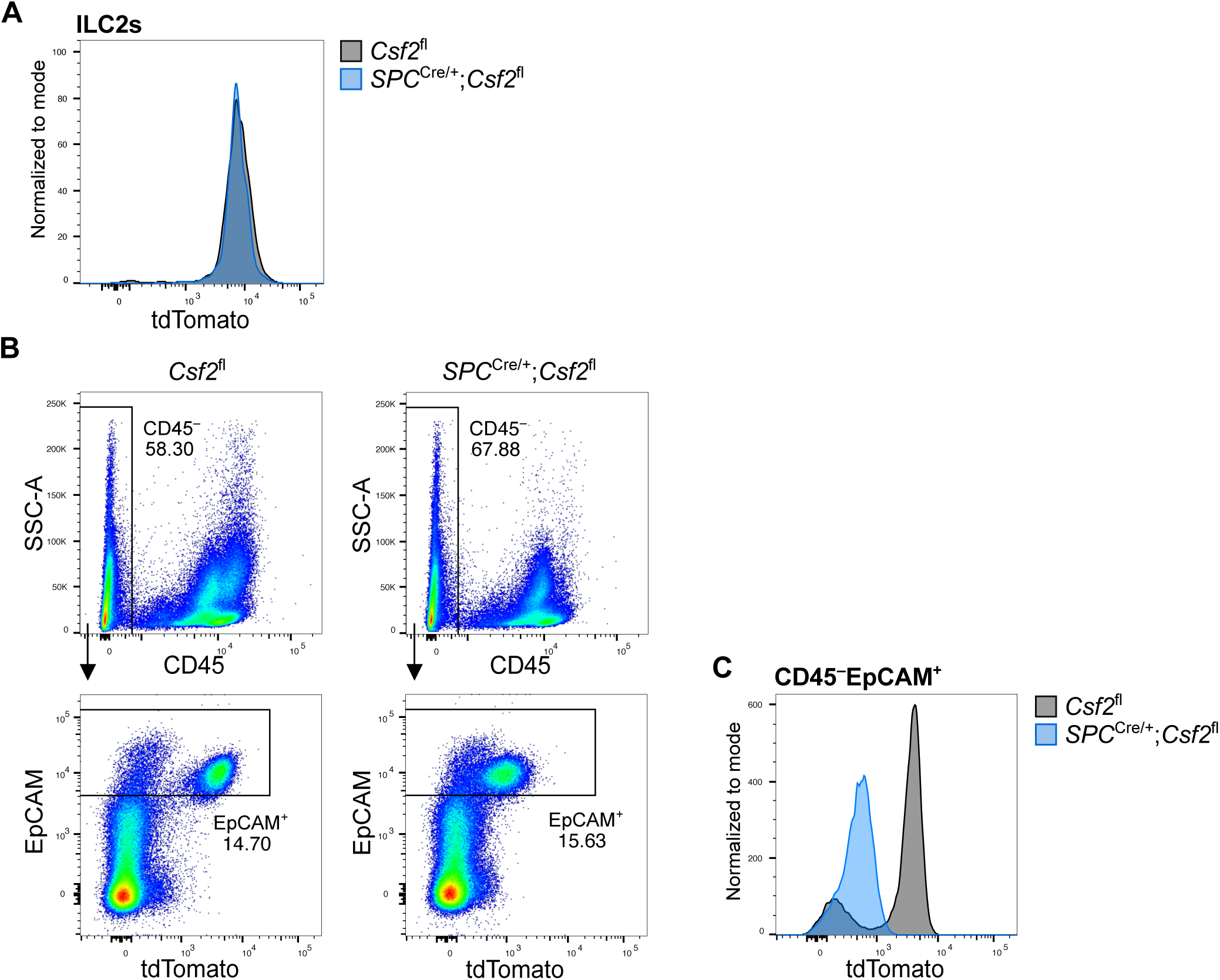
GM-CSF contribution of ILC2s and EpCAM^+^ epithelial cells in neonatal *SPC*^Cre^ mice. (**A**) Expression level of tdTomato in CD45^+^CD3^−^CD19^−^DX5^−^Lin^−^Thy1.2^+^ST2^+^ ILC2s from *Csf2*^fl^ (grey) and *SPC*^Cre/+^;*Csf2*^fl^ (blue) mice. (**B**) Gating strategy used to identify CD45^−^EpCAM^+^ cells. (**C**) Expression level of tdTomato in CD45^−^EpCAM^+^ epithelial cells from *Csf2*^fl^ (grey) and *SPC*^Cre/+^;*Csf2*^fl^ (blue) mice. (**A-C**) Data are from one experiment, representative of five independent experiments.

**Figure S4.**
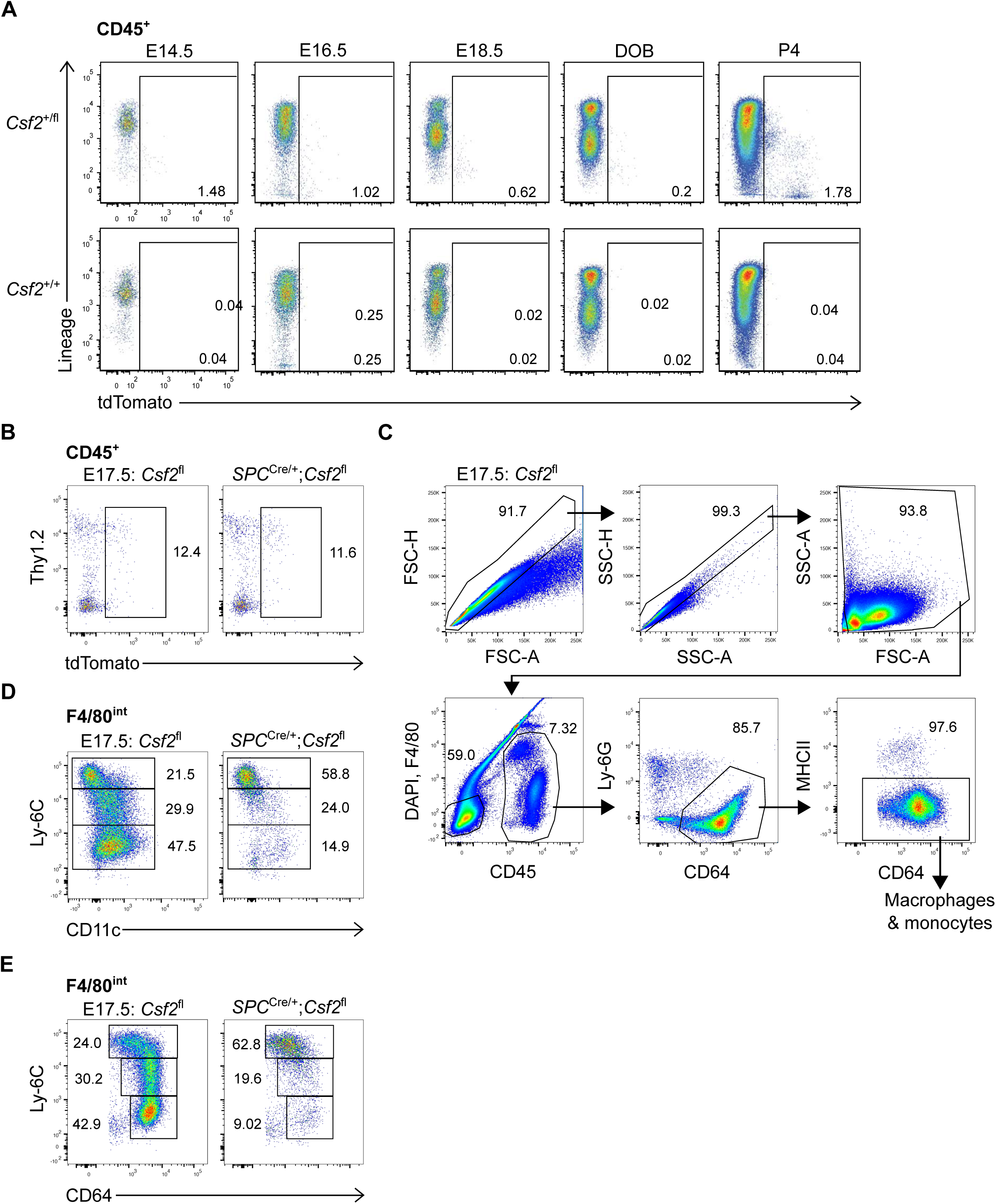
Kinetics of hematopoietic-derived GM-CSF during embryogenesis. **(A)** Kinetics of tdTomato expression in the CD45^+^ compartment of perinatal lungs of *Csf2*^+/fl^ and *Csf2*^+/+^ mice ranging from E14.5 to P4, determined by flow cytometry. (**B**) Flow cytometry analysis of CD45^+^ tdTomato^+^ populations in E17.5 lungs of *Csf2*^fl^ and *SPC*^Cre/+^;*Csf2*^fl^ mice. (**C**) Gating strategy used to identify primitive macrophages (G1 in Figure 5D) and fetal monocytes (G2 in Figure 5D). (**D**) Flow cytometry analysis of Ly-6C and CD11c levels in the developing fetal monocytes population (G2 in Figure 5D) in E17.5 lungs of *Csf2*^fl^ and *SPC*^Cre/+^;*Csf2*^fl^ mice. (**E**) Flow cytometry analysis of Ly-6C and CD64 levels in the developing fetal monocytes population (G2 in Figure 5D) in E17.5 lungs of *Csf2*^fl^ and *SPC*^Cre/+^;*Csf2*^fl^ mice. (**A**) Data representative of at least two independent experiments per time point. (**B-E**) Data are from one experiment, representative of three independent experiments.

**Figure S5.**
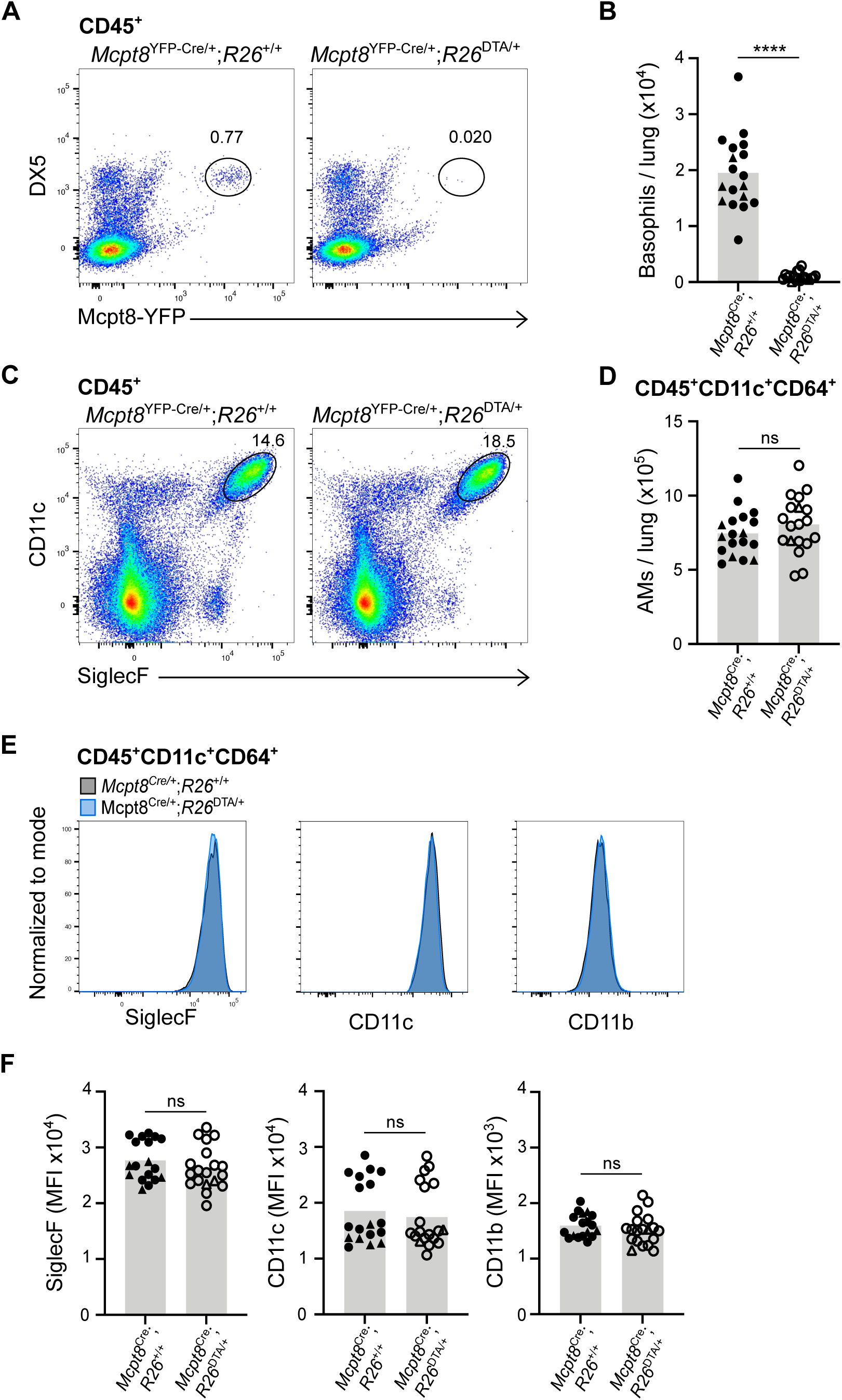
Constitutive depletion of basophils does not significantly alter AM populations in the adult lung. (**A**) Flow cytometry analysis CD45^+^DX5^+^Mcpt8-YFP^+^ basophils in adult lungs of *Mcpt8*^YFP-Cre^;R26^+/+^ and *Mcpt8*^YFP-Cre^;R26^DTA/+^ mice. (**B**) Quantification of CD45^+^DX5^+^Mcpt8-YFP^+^ basophils in adult lungs of *Mcpt8*^YFP-Cre^;R26^+/+^ and *Mcpt8*^YFP-Cre^;*R26*^DTA/+^ mice. *Mcpt8*^YFP-Cre/+^ mice are indicated by circles, while *Mcpt8*^YFP-Cre/YFP-Cre^ mice are indicated by triangles. (**C**) Flow cytometry analysis of CD45^+^CD11c^+^SiglecF^+^ AMs in adult lungs of *Mcpt8*^YFP-Cre^;*R26*^+/+^ and *Mcpt8*^YFP-Cre^;*R26*^DTA/+^ mice. (**D**) Quantification of AMs (CD45^+^CD11c^+^CD64^+^) in adult lungs of *Mcpt8*^YFP-Cre^;*R26*^+/+^ and *Mcpt8*^YFP-Cre^;*R26*^DTA/+^ mice. Genotypes are indicated as per (B). (**E**) Expression levels of SiglecF, CD11c and CD11b for AMs (CD45^+^CD11c^+^CD64^+^), as determined by flow cytometry, for adult *Mcpt8*^YFP-Cre^;*R26*^+/+^ (grey) and *Mcpt8*^YFP-Cre^;*R26*^DTA/+^ (blue) mice. (**F**) Quantification of SiglecF, CD11c and CD11b for AMs (CD45^+^CD11c^+^CD64^+^) in adult *Mcpt8*^YFP-Cre^;*R26*^+/+^ and *Mcpt8*^YFP-Cre^;*R26*^DTA/+^ mice. Genotypes are indicated as per (B). (**A, C, E**) Data are from one experiment, representative of three independent experiments. (**B, D, F**) Data are pooled from three independent experiments.

